# Intra- and inter-specific density dependence of body condition, growth, and habitat temperature in chub mackerel (*Scomber japonicus*)

**DOI:** 10.1101/2021.03.25.436928

**Authors:** Yasuhiro Kamimura, Makoto Taga, Ryuji Yukami, Chikako Watanabe, Sho Furuichi

**Author notes:** **Names and addresses of authors**: Y. Kamimura, R. Yukami, C Watanabe, and S. Furuichi. Fisheries Research Center, National Research Institute of Fisheries Science, Japan Fisheries Research and Education Agency, 2-12-4 Fukuura, Kanazawa, Yokohama, Kanagawa 236-8648, Japan M. Taga. Ibaraki Prefectural Fisheries Research Institute, 3551-8, Mitsuzuka, Hiraiso, Hitachinaka, Ibaraki 311-1203, Japan. **Corresponding author**: Y. Kamimura.

## Abstract

The density dependence of growth and body condition have important impacts on fish population dynamics and fisheries management. Although population density is known to affect the temperature of the habitat selected, how this affects the density dependence of growth and body condition remains unclear. Here, we investigated annual changes in body condition, habitat temperature, and cohort-specific growth of chub mackerel (*Scomber japonicus*) in the western North Pacific and examined quarterly changes in the density dependence of body condition. We hypothesized that chub mackerel body condition is affected both directly (e.g., through competition for food) and indirectly (through changes in habitat temperature) by the abundance of both conspecifics (i.e., chub mackerel) and heterospecifics (the Japanese sardine *Sardinops melanostictus*). Indeed, chub mackerel body condition, habitat temperature, and growth all decreased with increasing conspecific and heterospecific abundance. Mean annual growth rates in chub mackerel were positively corelated with body condition. The best model showed that conspecific and/or heterospecific abundance had strong negative effects on chub mackerel body condition in all seasons, and influenced habitat temperature in some seasons. By contrast, temperature effects on body condition were weak. Therefore, direct effects likely have more impact than indirect effects on density-dependent body condition and growth.

## Introduction

Density-dependent processes are fundamental to fish population dynamics (Rose *et al.*, 2001; Andersen *et al.*, 2017). Growth, survival, movement, and reproduction are all affected by population density, and can contribute to a decrease or increase in population growth rate at high or low densities (Rose *et al.*, 2001). In general, density-dependent mortality is a dominant factor in fish population dynamics in the early juvenile stage, and density-dependent growth becomes dominant in the late juvenile and adult stages (Lorenzen and Enberg, 2002; Lorenzen, 2008; Van Gemert and Andersen, 2018; Zimmermann *et al.*, 2018). Although the density dependence of recruitment is a stronger regulator of fish population size, the density dependence of growth affects productivity and catch composition and may be particularly important for fisheries management (Zimmermann *et al.*, 2018). In addition, rapid changes in fish population density and the subsequent drastic changes in growth and maturation often complicate stock assessment and fisheries forecasting (Eero *et al.*, 2015). There is an urgent need to understand the density dependence of growth to inform effective management decisions.

As with growth, body condition is often used as a proxy of individual fitness, as it reflects the size of individual energy reserves. Many studies have sought to identify determinants of body condition, such as conspecific and heterospecific density, temperature, prey availability, and fishing impacts, as a way to understand fish population dynamics (Rueda *et al.*, 2015; Brosset *et al.*, 2017). It is widely accepted that fish body condition can affect growth, reproductive potential, and survival (Blackwell *et al.*, 2000; Lloret *et al.*, 2002) and is an important data point for stock assessment and fisheries management (Blackwell *et al.*, 2000; Dahl *et al.*, 2019).

Temperature is one of the most important abiotic factors for fish growth and body condition (Houde, 1989; Rueda *et al.*, 2015). Many studies have considered temperature to be a density-independent factor when examining its effects on growth and body condition (Brunel and Dickey-Collas, 2010; Casini *et al.*, 2011, 2016; Brosset *et al.*, 2015; Cormon *et al.*, 2016). However, because the temperature of habitats selected by fish can vary with population density (Swain and Kramer, 1995), population density can indirectly affect growth and body condition through changes in habitat temperature. In addition, this indirect effect can vary with season. Comprehensive analyses of the effect of density-dependent changes in habitat temperature on growth and body condition, and how these effects change over the seasons, are required to properly understand the process of density dependence.

Chub mackerel (*Scomber japonicus*) is widely distributed across the North Pacific and is one of the most productive and economically important fisheries resources in the world, with annual catches reaching ca. 1.5 million metric tons (FAO, 2020). However, the abundance of the Pacific stock of chub mackerel declined in the 1990s and early 2000s (Yukami *et al.*, 2020a). Since 2013, when a strong year class occurred, its abundance has dramatically increased (Table 1). Although previous studies reported changes in factors such as growth and maturity during the period of population decline (Watanabe and Yatsu, 2004, 2006), limited information is available for the period of population increase, especially for recent years. In addition, although much attention has been paid to the impact of environmental and endogenous factors (e.g., temperature, food availability, and age/size) on chub mackerel growth in the larval and early juvenile stages (Hunter and Kimbrell, 1980; Sassa and Tsukamoto, 2010; Kamimura *et al.*, 2015; Higuchi *et al.*, 2019; Kaneko *et al.*, 2019; Taga *et al.*, 2019; Nakamura *et al.*, 2020), comparatively little research has been conducted on later growth stages (Parrish and Mallicoate, 1995; Watanabe and Yatsu, 2004; Yatsu *et al.*, 2019).

**Table 1.**
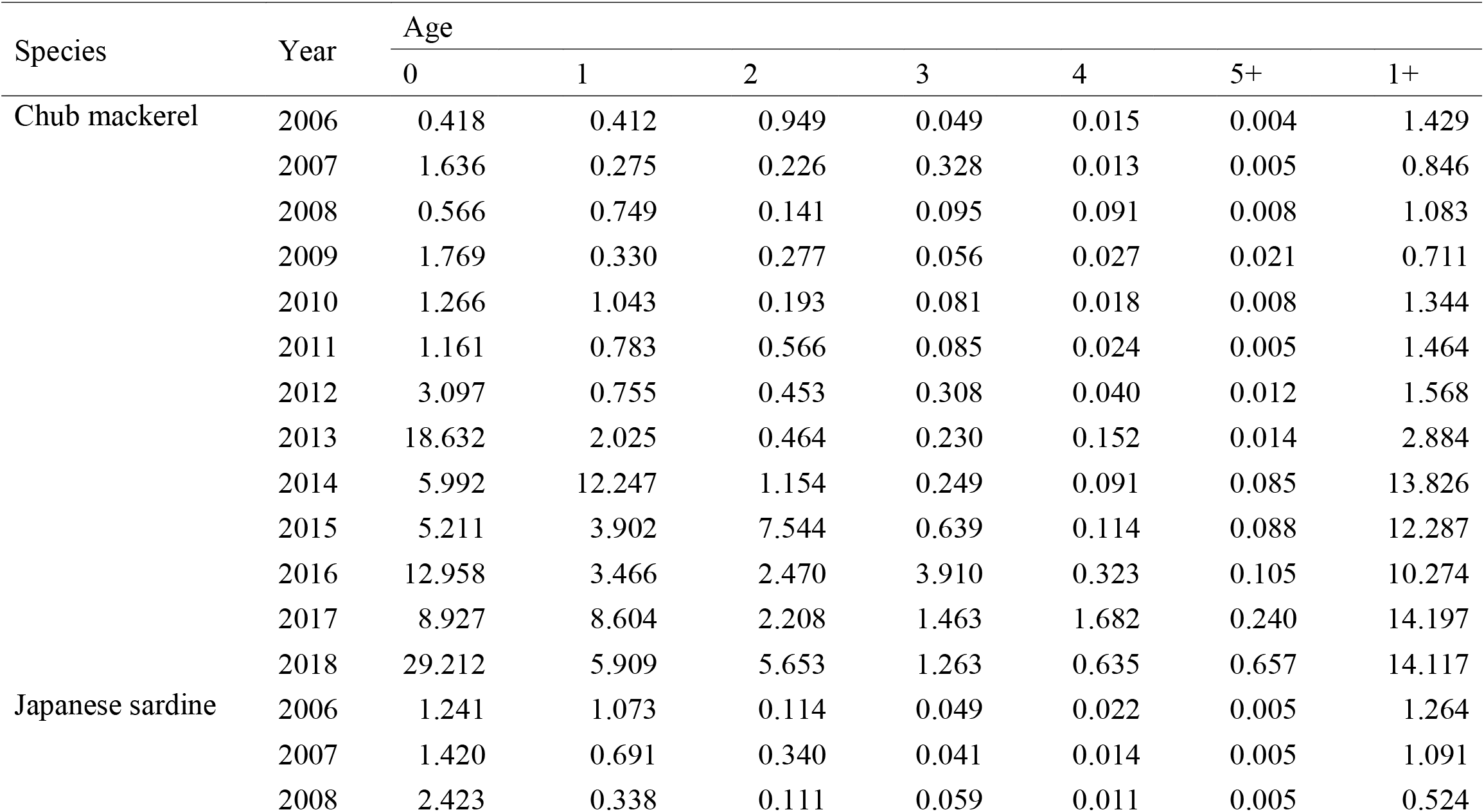
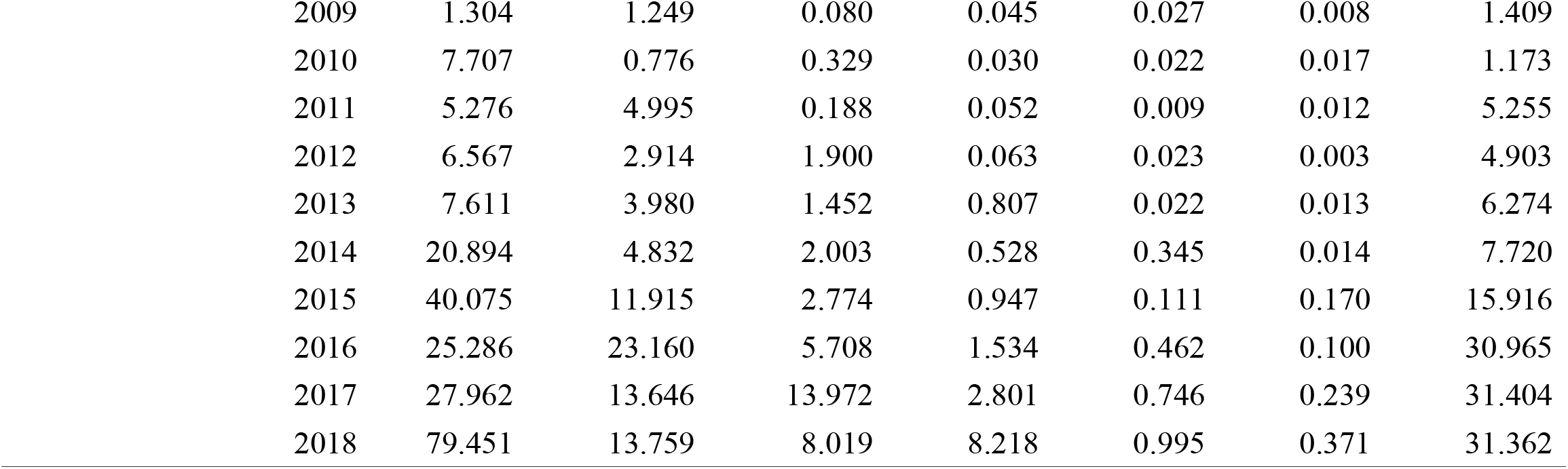
Abundances (10^9^ individuals) of the Pacific stock of chub mackerel and Japanese sardine as estimated by Yukami *et al.* (2020a) and Furuichi et al. (2020).

| Species | Year | Age |  |  |  |  |  |  |
| --- | --- | --- | --- | --- | --- | --- | --- | --- |
|  |  | 0 | 1 | 2 | 3 | 4 | 5+ | 1+ |
| Chub mackerel | 2006 | 0.418 | 0.412 | 0.949 | 0.049 | 0.015 | 0.004 | 1.429 |
|  | 2007 | 1.636 | 0.275 | 0.226 | 0.328 | 0.013 | 0.005 | 0.846 |
|  | 2008 | 0.566 | 0.749 | 0.141 | 0.095 | 0.091 | 0.008 | 1.083 |
|  | 2009 | 1.769 | 0.330 | 0.277 | 0.056 | 0.027 | 0.021 | 0.711 |
|  | 2010 | 1.266 | 1.043 | 0.193 | 0.081 | 0.018 | 0.008 | 1.344 |
|  | 2011 | 1.161 | 0.783 | 0.566 | 0.085 | 0.024 | 0.005 | 1.464 |
|  | 2012 | 3.097 | 0.755 | 0.453 | 0.308 | 0.040 | 0.012 | 1.568 |
|  | 2013 | 18.632 | 2.025 | 0.464 | 0.230 | 0.152 | 0.014 | 2.884 |
|  | 2014 | 5.992 | 12.247 | 1.154 | 0.249 | 0.091 | 0.085 | 13.826 |
|  | 2015 | 5.211 | 3.902 | 7.544 | 0.639 | 0.114 | 0.088 | 12.287 |
|  | 2016 | 12.958 | 3.466 | 2.470 | 3.910 | 0.323 | 0.105 | 10.274 |
|  | 2017 | 8.927 | 8.604 | 2.208 | 1.463 | 1.682 | 0.240 | 14.197 |
|  | 2018 | 29.212 | 5.909 | 5.653 | 1.263 | 0.635 | 0.657 | 14.117 |
| Japanese sardine | 2006 | 1.241 | 1.073 | 0.114 | 0.049 | 0.022 | 0.005 | 1.264 |
|  | 2007 | 1.420 | 0.691 | 0.340 | 0.041 | 0.014 | 0.005 | 1.091 |
|  | 2008 | 2.423 | 0.338 | 0.111 | 0.059 | 0.011 | 0.005 | 0.524 |

|  |  |  |  |  |  |  |  |
| --- | --- | --- | --- | --- | --- | --- | --- |
| 2009 | 1.304 | 1.249 | 0.080 | 0.045 | 0.027 | 0.008 | 1.409 |
| 2010 | 7.707 | 0.776 | 0.329 | 0.030 | 0.022 | 0.017 | 1.173 |
| 2011 | 5.276 | 4.995 | 0.188 | 0.052 | 0.009 | 0.012 | 5.255 |
| 2012 | 6.567 | 2.914 | 1.900 | 0.063 | 0.023 | 0.003 | 4.903 |
| 2013 | 7.611 | 3.980 | 1.452 | 0.807 | 0.022 | 0.013 | 6.274 |
| 2014 | 20.894 | 4.832 | 2.003 | 0.528 | 0.345 | 0.014 | 7.720 |
| 2015 | 40.075 | 11.915 | 2.774 | 0.947 | 0.111 | 0.170 | 15.916 |
| 2016 | 25.286 | 23.160 | 5.708 | 1.534 | 0.462 | 0.100 | 30.965 |
| 2017 | 27.962 | 13.646 | 13.972 | 2.801 | 0.746 | 0.239 | 31.404 |
| 2018 | 79.451 | 13.759 | 8.019 | 8.218 | 0.995 | 0.371 | 31.362 |

Chub mackerel mainly spawns in the waters around the Izu islands and the Boso peninsula, Japan (Figure 1a) from March to June (Watanabe and Yatsu, 2006; Kanamori *et al.*, 2019). Larvae and early juveniles are transported offshore from April to September by the Kuroshio and Kuroshio extension to a large region in the western North Pacific stretching from the Kuroshio–Oyashio transition region to the Oyashio region, after which juveniles begin to migrate to the coastal waters of northern Japan (Watanabe and Nishida, 2002; Takahashi *et al.*, 2014; Kamimura *et al.*, 2015). Age-1+ mackerel typically migrate northward along the Japanese coast from July to September and southward from October to December (Watanabe, 1970; Usami, 1973).

**Figure 1.**
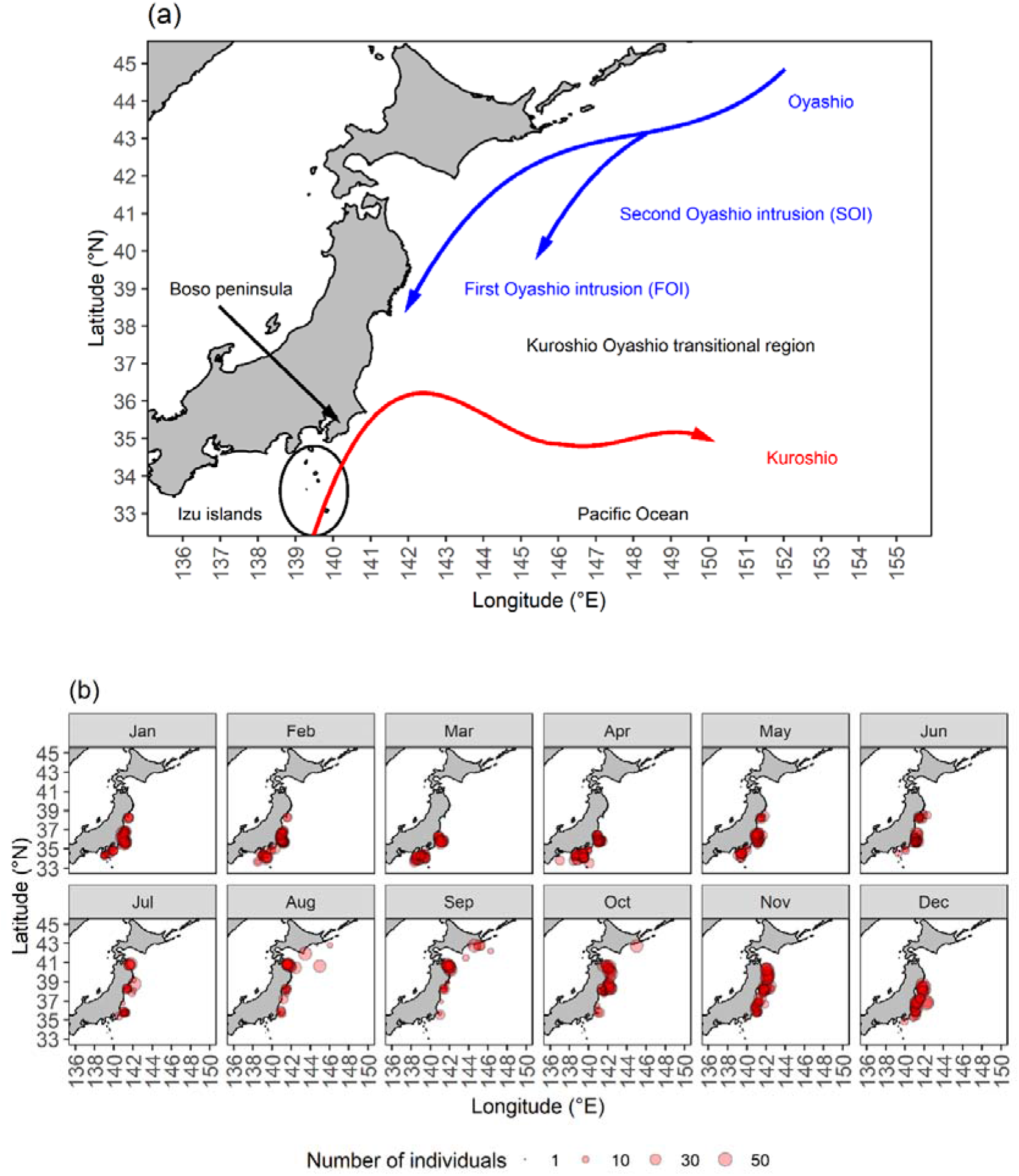
Maps of (a) the oceanographic features of the western North Pacific and (b) the monthly catch locations of chub mackerel used in this study.

Japanese sardine (*Sardinops melanostictus*) is also an important fishery resource in Japan, and the size of the Pacific stock has been increasing over the past ten years (Table 1). Seasonal changes in its distribution are similar to those of chub mackerel (Watanabe and Nishida, 2002; Yatsu *et al.*, 2005; Yatsu, 2019). The major prey items of juveniles and adults, such as Copepoda, Eupausiidae, Decapoda, and Amphipoda (Yamashita, 1955; Nakai, 1962; Kawasaki, 1984; Yamamoto and Katayama, 2012), overlap with those of late-juvenile and adult chub mackerel (Sato *et al.*, 1975; Nakatsuka *et al.*, 2010; Taga and Yamashita, 2018). Therefore, Japanese sardine is a potential competitor of chub mackerel (Kishida and Matsuda, 1993; Yatsu *et al.*, 2005), and a previous study has indicated that Japanese sardine density has a negative effect on chub mackerel recruitment (Kishida and Matsuda, 1993). However, the mechanisms of inter-specific density dependence of chub mackerel growth and body condition remain unknown.

The principal aim of this study was to clarify the density dependence of growth and body condition of late-juvenile and adult chub mackerel in the Pacific by considering the direct (e.g., through competition for food) and indirect (i.e., through changes in habitat temperature) effects of conspecific and heterospecific density. In addition, we also aimed to identify seasonal changes in density dependence and the drivers of fluctuations in body condition. First, individual size-at-age, relative condition factor, and habitat temperature were estimated to analyze the relationship between mean annual growth rate and mean body condition on a quarterly basis. Second, we used structural equation models (SEMs) to examine the effects of chub mackerel and Japanese sardine abundance, environmental conditions (i.e., the location of the Oyashio Current and the locations of sample capture), and an endogenous factor (age) on body condition and habitat temperature by quarter.

## Materials and methods

### Fish sampling and laboratory procedures

A total of 18,302 chub mackerel landed in fishing ports on the Pacific coast of northern Japan between 2006 and 2019 were used for the study (Figure 1b). Fishing gear included purse seines, dipnets, setnets, and vertical long lines. Most of the samples (12,379 individuals, or 68%) were captured by purse seine fisheries. Fork length (FL) was measured to the nearest 1 mm by using digital calipers. Total weight (TW), gonad weight (GW), and stomach content weight (SCW) were measured to the nearest 0.1 g by using digital scales. Sex was determined by direct gonad observation.

Ages were determined by counting annual rings on scales (Kondo and Kuroda, 1966) or sectioned sagittal otoliths (Kawashima *et al.*, 2017) under electronic microscopes. Previous studies have used gonad observation of mature fish and otolith microstructure analysis of juveniles to reveal that spawning mainly occurs in April (Watanabe, 2010; Kamimura *et al.*, 2015). Therefore, we assumed that the hatch date was in April and calculated ages as *t = A + m*/12, where *A* is age (in years) estimated from scale and otolith observations, and *m* is the number of months between the catch month and the preceding April.

### Growth analysis

A total of 15,415 fish in the 2006 to 2016 year classes were used for growth analysis. (Table 2). The von Bertalanffy (VB) equation was fitted separately to the FL-at-age data for each year class to estimate cohort-specific growth parameters:

**Table 2.** Fork length (FL) range, age (*t*) range, and estimated von Bertalanffy parameters (*L*_inf_, *K*, and *t*_0_) by year class for chub mackerel.

| Year<br>class | $n$ | FL range<br>(mm) | $t$ range<br>(years) | $L_{inf}$ | $K$ | $t_0$ |
| --- | --- | --- | --- | --- | --- | --- |
| All | 15 415 | 121–470 | 0.2–11.1 | 371.0 | 0.39 | –1.96 |
| 2006 | 525 | 207–426 | 0.4–8.1 | 440.5 | 0.26 | –2.25 |
| 2007 | 1 548 | 188–460 | 0.3–9.8 | 400.4 | 0.49 | –1.26 |
| 2008 | 659 | 226–434 | 0.6–11.1 | 396.0 | 0.50 | –1.38 |
| 2009 | 1 315 | 212–440 | 0.8–10.1 | 400.6 | 0.44 | –1.62 |
| 2010 | 1 351 | 208–470 | 0.4–9.3 | 396.5 | 0.48 | –1.59 |
| 2011 | 830 | 226–455 | 0.8–8.3 | 393.5 | 0.35 | –2.88 |
| 2012 | 1 865 | 121–457 | 0.3–7.3 | 360.2 | 0.55 | –1.41 |
| 2013 | 4 522 | 160–460 | 0.5–6.7 | 356.9 | 0.32 | –2.41 |
| 2014 | 1 533 | 198–430 | 0.4–5.7 | 348.4 | 0.36 | –2.37 |
| 2015 | 647 | 193–420 | 0.4–4.7 | 400.5 | 0.25 | –2.23 |
| 2016 | 620 | 136–358 | 0.2–3.7 | 339.9 | 0.47 | –1.29 |

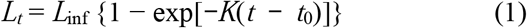

where *L*_*t*_, *L*_inf_, *K*, and *t*_0_ are FL at age *t*, asymptotic FL, growth coefficient, and hypothetical age at zero FL, respectively. Non-linear least-squares regression was used to estimate these parameters. Initial values were estimated by using the Ford–Walford plot. Sex-specific models were calculated for each year class, and VB growth formulae were compared between males and females within each year class by using the *F* test.

Annual growth rates in length (*G*_*L*_) for ages 1–5 were calculated as follows:

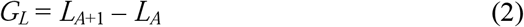

where *L*_*A*+1_ is the FL of chub mackerel at age *A* + 1, and *L*_*A*_ is the age *A* estimated by the VB growth formula for each year class.

### Body condition

Relative condition factor (*K_n_*; Le cren, 1951; Froese, 2006) was computed to examine individual fish condition by using 8429 age-1+ chub mackerel from the 2006–2016 year classes (caught from 2007 to 2018 in 510 hauls) whose catch locations were recorded.

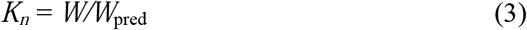

where *W* is body weight (excluding GW and SCW) and *W*_pred_ is the predicted *W* from a weight–length relationship of the form:

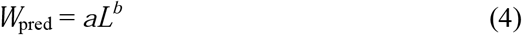

where *L* is FL, *a* is the allometric coefficient between weight and length, and *b* is the exponent of that relationship. We estimated these parameters by using non-linear least-squares regression.

### Abundance indices of the Pacific stock of chub mackerel and Japanese sardine

Abundances of the Pacific stock of chub mackerel and Japanese sardine were estimated by virtual population analysis from the 1970s to 2018 (Furuichi *et al.*, 2020; Yukami *et al.*, 2020a). We used these estimates as abundance indices for this study. As mentioned above, the distributions of age-0 chub mackerel and Japanese sardine are different from those of older individuals; hence, the age-1+ abundances were used as the abundance indices. Age-1+ chub mackerel abundances (*N*_*m*_) ranged from 8.5 × 10^9^ (2007) to 141.2 × 10^9^ individuals (2017), and Japanese sardine abundances (*N*_*s*_) ranged from 5.2 × 10^9^ (2008) to 313.6 × 10^9^ individuals (2018; Table 1). *N*_*m*_ increased dramatically in 2014 and reached about 140 × 10^9^ individuals. *N*_*s*_ gradually increased after 2011 and stabilized at about 300 × 10^9^ individuals after 2016.

### Confirmation of overlapping distributions of chub mackerel and Japanese sardine by comparing catch locations

To confirm the distributional overlap of chub mackerel and Japanese sardine in recent years, we compared mackerel (chub and spotted mackerel, *S. australasicus,* were combined) and Japanese sardine catch locations by season by using catch reports of purse seine fisheries made by the Japan Fisheries Agency from 2010 to 2017.

### Habitat temperature

Seasonal changes in the vertical distribution of chub mackerel were reported in previous studies (Usami, 1973; Sato, 1974). Hence, we examined mean monthly habitat depths based on 266 school-depth records from fishing vessel logbooks recorded by the North Pacific district purse seine fishery for 2008–2019. Mean monthly school depths showed that mackerel habitat was deeper in winter (November to February) than in summer (May to August; Figure 2).

**Figure 2.**
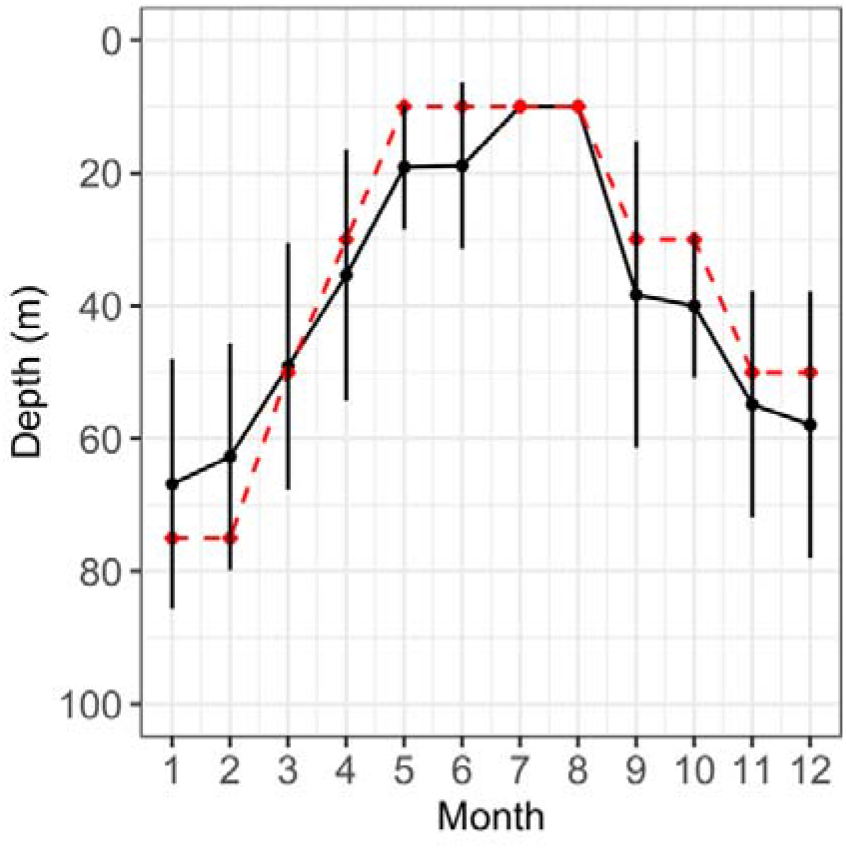
Monthly changes in chub mackerel vertical distribution. The black solid line indicates the mean school depth recorded in logbooks by purse seine fisheries, and the red dashed line indicates the habitat depths used when estimating habitat temperatures with FRA-ROMS. Vertical error bars show standard deviations.

We used the temperatures at which chub mackerel had been caught (TC) as a proxy for habitat temperature in this study. Temperature data were obtained from reanalysis data of the Regional Ocean Modelling System (ROMS) developed by the Japan Fisheries and Education Agency (FRA) for fisheries science (FRA-ROMS; Kuroda *et al.*, 2017). Temperatures were estimated by using fishing locations and dates recorded in the logbooks. FRA-ROMS provides averaged daily temperatures at 1/10-degree scale at depths of 0, 10, 30, 50, 75, 100, and 200 m in the Kuroshio–Oyashio region. Therefore, we extracted temperatures for the depths provided by FRA-ROMS that were closest to the mean monthly school depths estimated above (i.e., 75 m in January and February; 50 m in March, November, and December; 30 m in April, September, and October; and 10 m in May, June, July, and August) (Figure 2).

### The First Oyashio Intrusion

The Oyashio Current is a western boundary current of western subpolar gyre in the North Pacific (Qiu, 2001) that transports cold, low-salinity subarctic water together with chemical and organic matter along the Kuril, Hokkaido, and Honshu (the main island in the Japanese archipelago) islands (Figure 1a). The southward current on the continental slope along the east coast of Honshu is referred to as the First Oyashio Intrusion (FOI), and plays an important role in the supply of nutrients and copepods to the region (Shimizu *et al.*, 2009; Kuroda *et al.*, 2017). Therefore, the FOI could potentially impact the growth and body condition of fisheries species. In this study, the southernmost position of the FOI (the SFOI) was determined from FRA-ROMS reanalysis data by using a procedure described in previous studies (Murakami, 1994; Kuroda *et al.*, 2017). We defined the SFOI as the point at which the southward convex locations of the 5-°C isotherm at a depth of 100 m (as calculated from approximately 10-d means) was the closest to the coast of Honshu. SFOIs were averaged for each month during 2006–2018.

### Statistical analysis

The piecewise SEM (Lefcheck, 2016) was used to investigate the effects of biotic and abiotic factors on the body condition of age-1+ chub mackerel. Classical SEMs assume that all observations are independent and that all variables are normally distributed. By contrast, in the piecewise SEM, the path diagram is translated to a set of linear equations and parameters are calculated by solving each equation separately; hence, the piecewise SEM can include random effects and distributions of various shapes (Lefcheck, 2016). We developed an *a priori* SEM (Figure S1) by using previous studies to parameterize the relationships between fish condition factor (*K*_*n*_), chub mackerel abundance (*N*_*m*_), Japanese sardine abundance (*N*_*s*_), age (*A*), the southernmost position of the FOI (SFOI), habitat temperature (TC), catch latitude (Lat), and catch longitude (Lon) (Parrish and Mallicoate, 1995; Watanabe and Yatsu, 2004). We hypothesized that these causal variables affect body condition indirectly through TC, as well as directly. Because monthly mackerel condition factor and temperature are most closely correlated at short lag times (i.e., a lag of 0 or 1 month) (Parrish and Mallicoate, 1995), TC was used as a causal variable for *K*_*n*_ in this study. Piecewise SEMs were constructed for each quarter starting in April, with the quarters defined as follows: Q1, April to June; Q2, July to September; Q3, October to December; and Q4, January to March. We assumed Q1 as the spawning season, Q2 as the season for the northward feeding migration, Q3 as the season for the southward migration, and Q4 as the pre-spawning season, based on previous studies of seasonal distribution (Watanabe, 1970; Usami, 1973) and reproductive surveys of chub mackerel (Watanabe and Yatsu, 2006). The goodness-of-fit of each model was evaluated using Shipley’s d-separation test (Shipley, 2000, 2009), which tests the assumption that all variables are conditionally independent, and we considered the final SEM consistent with the data if Fisher’s *C* statistic was *p* > 0.05. The model with the lowest Bayesian information criterion (BIC) was considered the final model; BIC is appropriate for confirmatory analyses and for finding the correct model (Aho *et al.*, 2014). Standardized path coefficient estimates were calculated for each path model.

To construct the piecewise SEM, the *K*_*n*_ model and TC model were separately fitted with individual regressions by using linear mixed models (LMMs). For the *K*_*n*_ model, *K*_*n*_ was used as a response variable; *N*_*m*_, *N*_*s*_, TC, *A*, SFOI, Lat, and Lon were used as explanatory variables; and sex (male, female, and unidentified) and haul ID (a categorical variable) were included as random variables. For the TC models, TC was used as a response variable; *N*_*m*_, *N*_*s*_, *A*_*t*_, SFOI, Lat, and Lon were used as explanatory variables; and sex was included as a random variable. The multicollinearity of models was checked by calculating a variance inflation factor for each explanatory variable. Collinearity was assessed with a cut-off value of 5 (James *et al.*, 2013); as a result, *N*_*s*_ and Lon were eliminated from the models in Q1 and Lon was eliminated in Q4.

To examine whether chub mackerel body condition was related to *G*_*L*_, relationships between mean *K*_*n*_ in each quarter and *G*_*L*_-at-age were analyzed by using LMMs. We used *G*_*L*_ for ages 1–5 as a response variable and mean *K*_*n*_ by age as explanatory variables with age as a random intercept. A likelihood-ratio test was conducted to examine the significance of *K*_*n*_ in this analysis.

All statistical tests were performed in R v. 3.6.3 (R Core Team, 2020). VB parameters were estimated by using the package *FSA* (Ogle *et al.*, 2020). Piecewise SEMs were constructed with the package *piecewiseSEM* (Lefcheck, 2016), and LMMs were analyzed with the package *lme4* (Bates *et al.*, 2015).

## Results

Chub mackerel was caught mainly from January to June in the waters offshore of the central Japanese archipelago between 34°N and 39°N (Figure 1b). During summer (July to September), catch locations were mainly in northern waters between 37°N and 43°N, and after October, they moved southward. The catch locations of mackerels (i.e., chub and spotted mackerel) and Japanese sardine in purse seine fisheries followed similar seasonal patterns and overlapped from 2010 to 2017 (Figure S2). We used catch data on chub mackerel (Yukami *et al.*, 2020a) and spotted mackerel (Yukami *et al.*, 2020b) to calculate the percentage of chub mackerel in the total mackerel catch in the Pacific stock. The percentages for each study year were 2010, 40.4%; 2011, 36.1%; 2012, 47.8%; 2013, 67.0%; 2014, 71.6%; 2015, 84.5%; 2016, 90.7%; and 2017, 93.6%. We estimated that most of the catches from 2010 to 2017 were in fact chub mackerel.

The mean monthly SFOI changed seasonally, with high latitudes in October–January followed by a rapid transition to low latitudes (Figure 3). The highest (farthest north) SFOI was observed in December 2015 (42°4’N), and the lowest (farthest south) in March 2017 (37°4’N).

**Figure 3.**
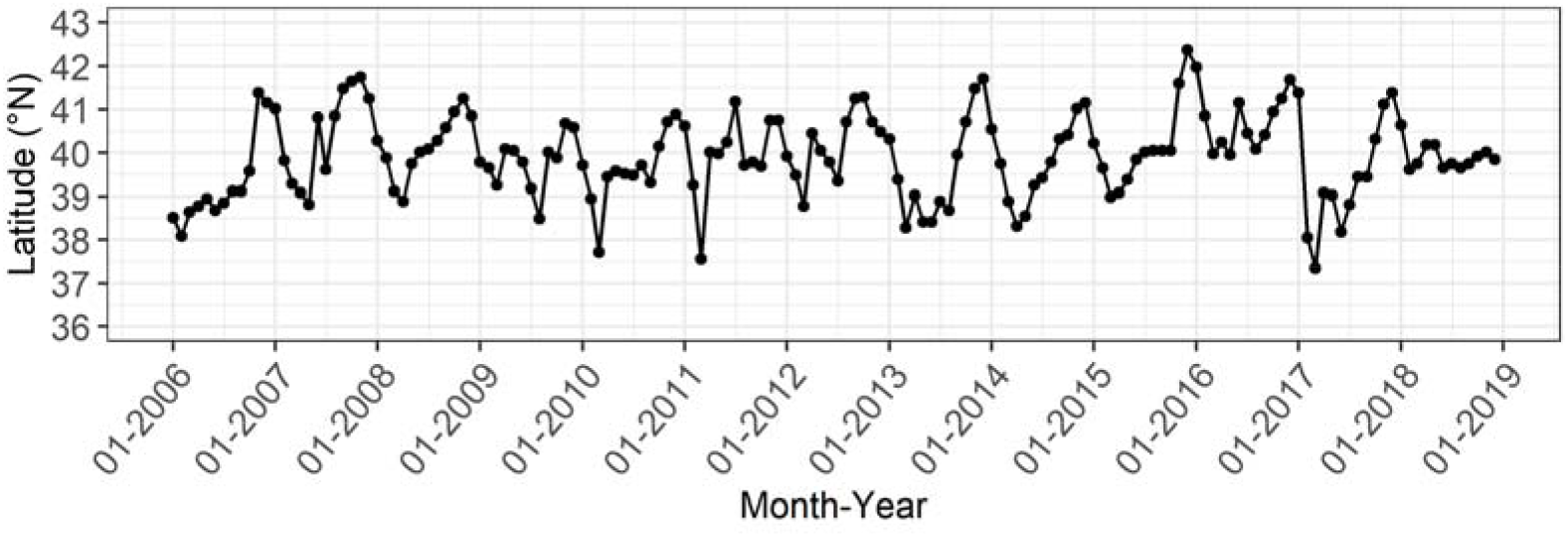
Monthly changes in the southernmost position of the First Oyashio Intrusion from January 2006 to December 2018.

The FLs of individuals used in the VB growth formulae ranged from 121 to 470 mm, and ages ranged from 0.2 to 11.1 years. VB growth formulae were fitted to each year class by sex. We found significant differences between male and female formulae in the 2008, 2009, and 2013 year classes (Figure S3 and Table S1). However, differences by sex in these year classes were small after age 1; therefore, we combined data from males, females, and individuals of unidentified sex to establish VB growth formulae for each year class.

Estimates of *L*_*inf*_, *K*, and *t*_0_ ranged from 339.9 (for the 2016 year class) to 440.5 (the 2006 year class), from 0.25 (for the 2015 year class) to 0.55 (the 2012 year class), and from −2.88 (for the 2011 year class) to −1.26 (the 2007 year class), respectively (Table 2). *G*_*L*_ ranges were 29.6–51.2 at age 1, 20.8–33.3 at age 2, 13.4–25.6 at age 3, 7.7–19.6 at age 4, and 4.4–15.1 at age 5 (Table S2). The estimated FLs were especially high for the 2007–2011 year classes and low for the 2013–2016 year classes, with the 2012 year class showing intermediate growth (Figure 4 and Table S2). The *G*_*L*_s of 2012 year class were lower after age 2 than the other year classes (Table S2).

**Figure 4.**
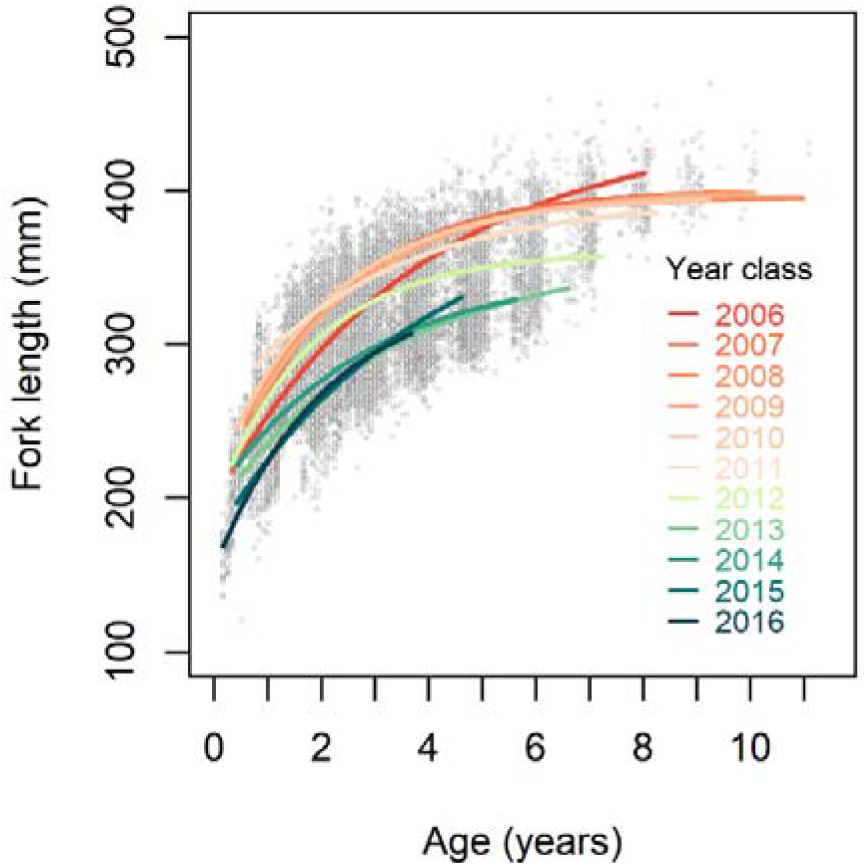
Comparison of von Bertalanffy growth formulas for chub mackerel among 11 year classes. Grey dots show the raw data, and coloured lines show von Bertalanffy curves fitted to each year class.

The weight–length relationship was estimated as *W* = 1.09 × 10^−6^*L*^3.41^. Values of *K*_*n*_ followed a generally downward trend in recent years and were lower in Q1 among older individuals and higher in Q3 and Q4 (Figure 5a and Tables S3 and S4). TC also declined over time and was higher among older individuals in Q1 and Q4. (Figure 5b and Tables S3 and S4). The mean TC in Q1 was relatively high (around 20 °C) until 2013, and fell to around 16 °C after 2014. Similarly, the mean TC in Q3 was around 20 °C until 2012, after which it fell to around 16 °C. The mean TC in Q4 dropped from around 17 °C to 14 °C beginning in 2014. The mean TC in Q2 ranged from 15 to 22 °C and there was no clear trend.

**Figure 5.**
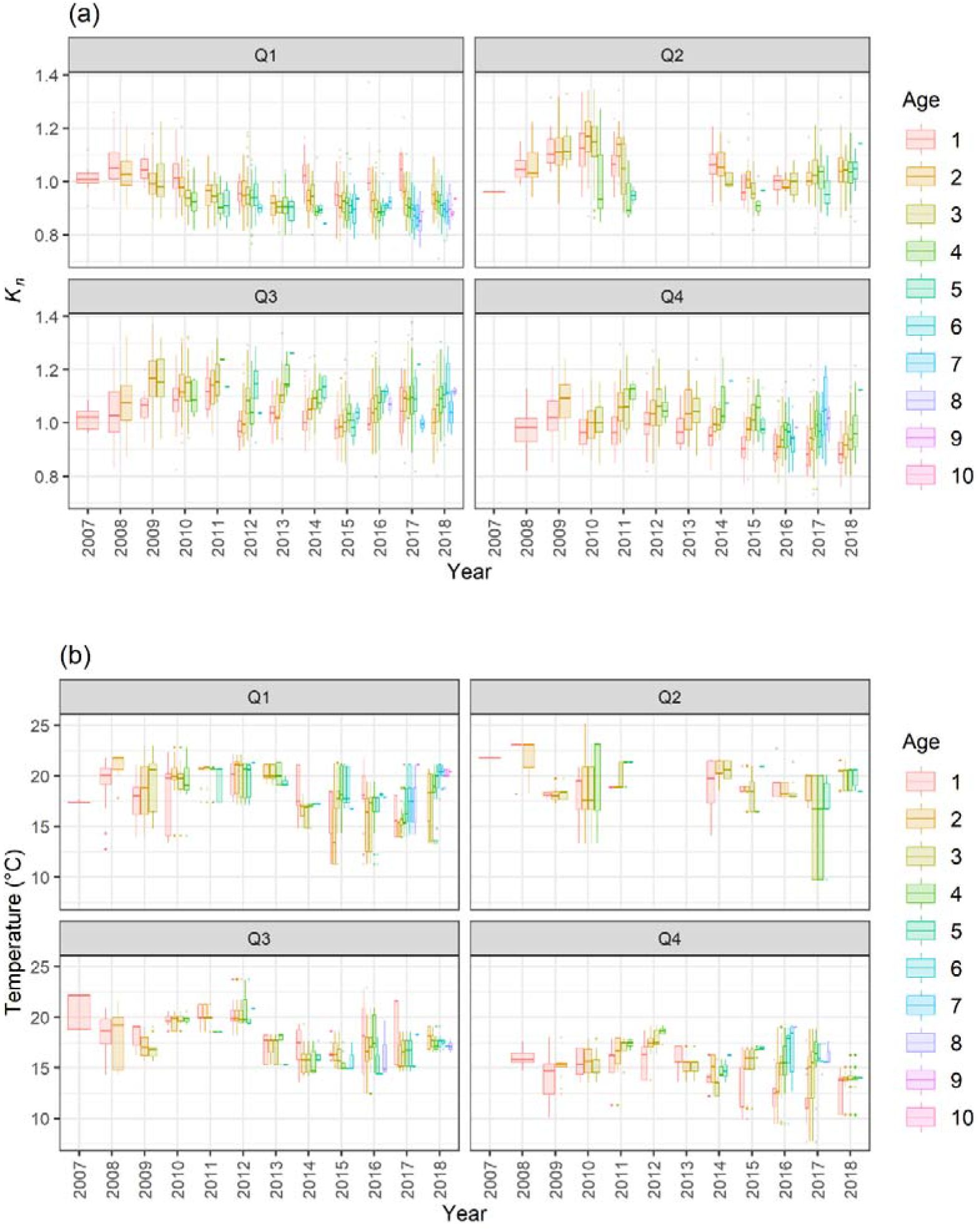
Box plots of (a) chub mackerel relative condition factor (*K*_*n*_) and (b) the estimated temperatures at which individuals were caught by age and quarter from 2006 to 2018.

*G*_*L*_ was significantly correlated with mean *K*_*n*_ in each quarter (Q1: χ^2^ = 34.6, *p* < 0.001; Q2: χ^2^ = 7.23, *p* = 0.007; Q3: χ^2^ = 6.50, *p* = 0.011; Q4: χ^2^ = 4.83, *p* = 0.028). Our model showed that *G*_*L*_ increased with higher *K*_*n*_ for ages 1–5 in all quarters (Figure 6 and Table 3).

**Figure 6.**
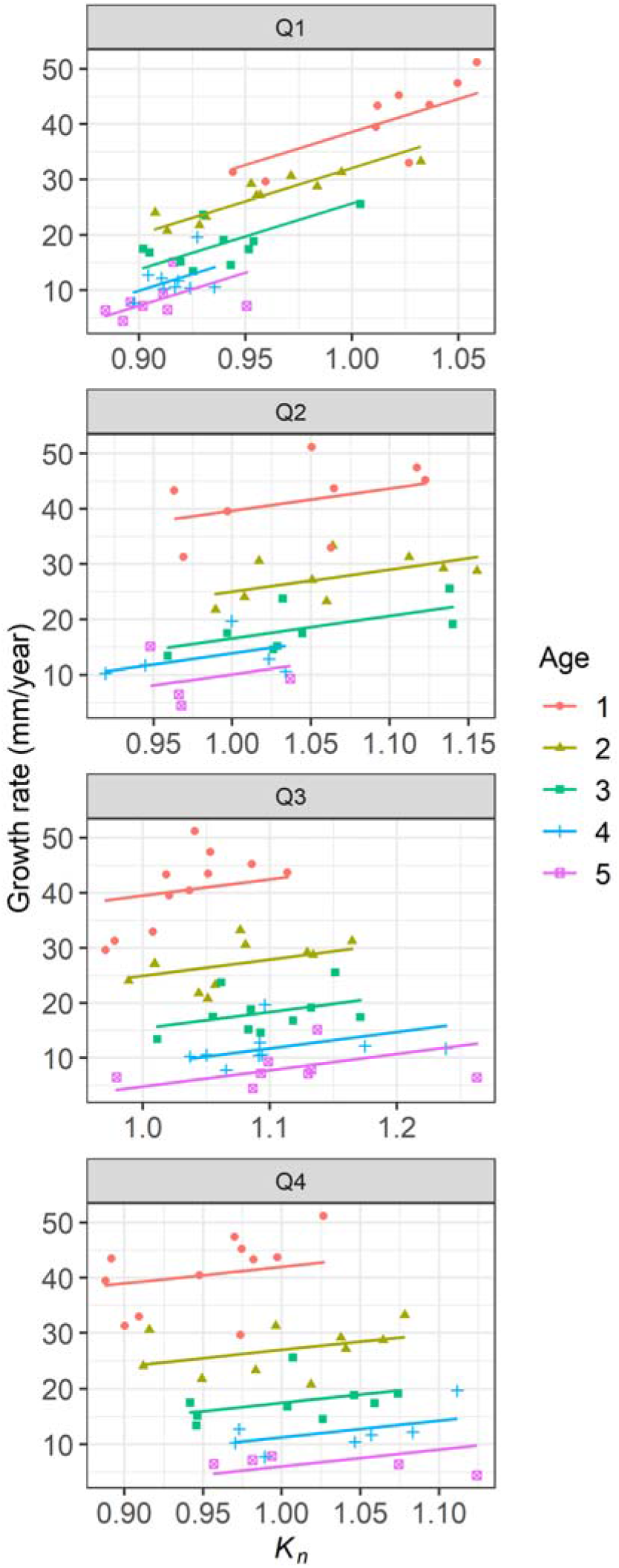
Relationships between mean relative condition factor (*K*_*n*_) and annual growth-at-age (*G*; mm year^−1^) by quarter. Lines represent linear functions estimated by using linear mixed models of *G*_*L*_ and *K*_*n*_.

**Table 3.** Parameter estimates of the models showing the effects of mean relative condition factor (*K*_*n*_) on growth rate in terms of length (*G*_*L*_) by quarter for chub mackerel.

| Quarter | Parameter | Estimate | SE |
| --- | --- | --- | --- |
| Q1 | Intercept | -91.84 | 16.06 |
| | $K_n$ | 119.32 | 16.63 |
| Q2 | Intercept | -19.35 | 15.35 |
| | $K_n$ | 40.39 | 14.23 |
| Q3 | Intercept | -11.01 | 13.27 |
| | $K_n$ | 29.69 | 11.09 |
| Q4 | Intercept | -9.34 | 14.32 |
| | $K_n$ | 30.08 | 13.06 |

The final SEM for each quarter was selected based on BIC (Table S5), and provided good fit to the data (Q1: Fisher’s *C* = 4.21, *p* = 0.38; Q2: Fisher’s *C* = 1.07, *p* = 0.98; Q3: Fisher’s *C* = 5.39, *p* = 0.25; and Q4: Fisher’s *C* = 6.14, *p* = 0.19). In these SEMs, biotic and abiotic effects on mackerel *K*_*n*_ and TC varied by quarter (Figure 7 and Table S6). Homogeneity tests of residuals for the best models showed that the residuals approximately follow a normal distribution, meaning that the assumption of homogeneous variances was met (Figures S4 and S5).

**Figure 7.**
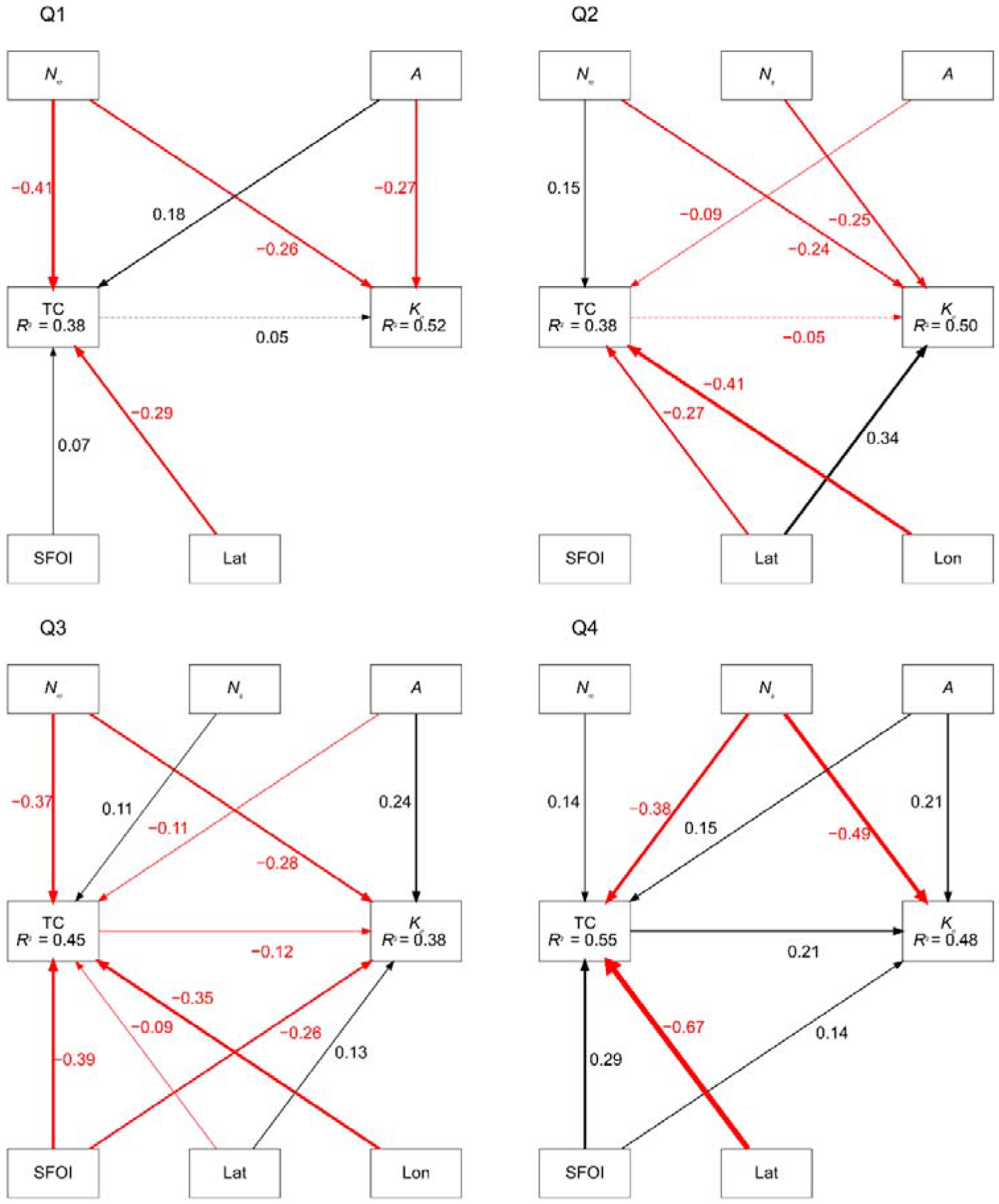
Final piecewise structural equation model of chub mackerel condition factor (*K*_*n*_) by quarter. The variables examined were chub mackerel abundance (*N*_*m*_), Japanese sardine abundance (*N*_*s*_), age (*A*), habitat temperature (TC), southernmost position of the First Oyashio Intrusion (SFOI), catch latitude (Lat), and catch longitude (Lon). Black arrows and red arrows represent positive and negative relationships, respectively. Solid lines indicate significant relationships (*p* < 0.05), and dotted lines indicate non-significant relationships. Standardized path coefficients are shown in black or red text, and the thickness of the paths correspond to the magnitude of these coefficients. *R*^2^ values inside the boxes are conditional *R*^2^.

*N*_*m*_ had strong negative direct effects on *K*_*n*_ in Q1, Q2, and Q3, and *N_s_* had strong negative direct effects in Q2 and Q4 (Figure 7 and Table S6). *A* had direct positive effects on *K_n_* in Q3 and Q4, but had a negative direct effect in Q1. SFOI had a negative direct effect on *K_n_* in Q3 and a positive direct effect in Q1 and Q4. Lat had positive direct effects on *K*_*n*_ in Q2 and Q3. TC had a negative effect on *K*_n_ in Q2 and Q3 and a positive effect in Q1 and Q4, but its effects were non-significant in Q1 and Q2.

*N*_*m*_ had positive effects on TC in Q2 and Q4 and negative effects in Q1 and Q3. *N_s_* had positive effects on TC in Q3 and a negative effect in Q4 (Figure 7 and Table S6). The indirect effects of *N_m_* on *K_n_* via TC were negative in Q1 (−0.41 ×0.05 = −0.02) and Q2 (0.15 × −0.05 = −0.01) and positive in Q3 (−0.37 × −0.12 = 0.04) and Q4 (0.14 × 0.22 = 0.03). The indirect effects of *N_s_* were negative in Q3 (0.11 × −0.12 = −0.01) and Q4 (−0.38 × 0.22 = −0.08). *A* had positive effects on TC in Q1 and Q4 and negative effects in Q2 and Q3, and had positive indirect effects on *K_n_* via TC in Q3 (−0.11 × −0.12 = 0.01) and Q4 (0.15 × 0.22 = 0.03). The effects of SFOI on *K_n_* differed among quarters, but had positive indirect effects on *K_n_* via TC in Q3 (−0.38 × −0.12 = 0.05) and Q4 (0.29 × 0.22 = 0.06). Although the direct and indirect effects of SFOI on *K_n_* in Q3 were in opposite directions, the direct effects of SFOI were of greater magnitude than the indirect effects. Lat and Lon had negative effects on TC in all quarters.

## Discussion

Over the past decade, chub mackerel body condition and growth have declined with increasing chub mackerel and Japanese sardine abundance in the western North Pacific (Figures 4 and 5a). This has likely been caused by the negative intra- and inter-specific density dependence of chub mackerel body condition. Positive relationships between body condition and mean annual growth rate in terms of length in fish aged 1–5 (Figure 6) suggest that body condition affects growth, and growth is affected by similar mechanisms as body condition. Previous studies on chub mackerel in the western North Pacific have underlined the importance of intra-specific density-dependent changes in growth, maturity, and egg production (Watanabe and Yatsu, 2004, 2006; Takasuka *et al.*, 2020). Our results show that chub mackerel body condition is as strongly affected by Japanese sardine abundance as by chub mackerel abundance. Therefore, we suggest that the inter-specific density dependence of body condition and growth are important factors for understanding chub mackerel population dynamics. Moreover, our data indicate that the carrying capacity of the western North Pacific for small pelagic fishes may have been reached within the last ten years.

In general, higher population densities cause lower food consumption and higher metabolic cost due to an increase in foraging effort, resulting in slower fish growth (Biro *et al.*, 2003; Amundsen *et al.*, 2007). In the case of chub mackerel, this appears to apply not only to the population density of chub mackerel itself but also to that of Japanese sardine. Chub mackerel and Japanese sardine are known to overlap in their feeding habits; additionally, in the present study, we show that the distribution of chub mackerel likely overlapped with that of Japanese sardine from 2010 to 2017 (Figure S2). Therefore, the decline in chub mackerel body condition associated with an increase in the density of conspecifics and heterospecifics was possibly mediated by changes in food consumption rate and/or metabolic cost. A previous study in Norwegian waters showed that the length- and weight-at-age of adult Atlantic mackerel (*Scomber scombrus*) decreases with increasing mackerel and Atlantic herring (*Clupea harengus*) density, suggesting that intra- and inter-specific food competition in summer feeding grounds is responsible for growth regulation (Olafsdottir *et al.*, 2016). In our study, our piecewise SEM analyses confirmed that both intra- and inter-specific density dependence negatively affected mackerel body condition in the northward feeding migration season (Figure 7 and Table S6), when chub mackerel and other small pelagic fishes generally converge on the same subarctic summer feeding grounds in the western North Pacific (Yatsu, 2019). Seasonal changes in the negative effects of chub mackere and Japanese sardine abundance on body condition, as estimated by our models, might indicate the timing of the occurrence of intra- and inter-specific competition. If the competition is over food resources, as is suggested in our study, future work that directly compares seasonal changes in the diets of chub mackerel and Japanese sardine could provide further insight into the possible mechanisms.

High body condition was associated with low temperatures during the southern migration and high temperatures in the pre-spawning season (Figure 7 and Table S6). Temperature likely affects fish metabolism directly as well as indirectly through the regulation of food resources (Lloret *et al.*, 2014), and the relationship between temperature and body condition is specific to species, seasons, and ecosystems. For example, one study reported a negative relationship between water temperature and chub mackerel body condition in the eastern North Pacific (Parrish and Mallicoate, 1995), and suggested that higher temperatures, which reflected a reduction in the flow of the California Current, likely caused a decrease in food availability. In contrast, temperature was positively related to the body condition of sprat (*Sprattus sprattus*), a thermophilic species in the Baltic Proper (Casini *et al.*, 2011). In our study area, because complex ocean structures are formed by the collision of the cold Oyashio and warm Kuroshio currents, there is a great deal of seasonal variability in productivity and fish food availability (Yasuda, 2003; Kobari *et al.*, 2004, 2008; Yatsu *et al.*, 2013; Itoh *et al.*, 2015), and the relationship between body condition and temperature may have differed among seasons.

Our results suggest that chub mackerel habitat temperatures were affected by intra- and inter-specific density dependence during the spawning, southward migration, and pre-spawning seasons in recent years. During the southward migration and pre-spawning seasons, Japanese sardine abundance had negative indirect effects (i.e., through habitat temperature) on chub mackerel body condition. This suggests that inter-specific competition displaced chub mackerel into habitats of different temperature, resulting in a decline in body condition. However, temperature effects on body condition are relatively small, and judging from the standardized path coefficients of the final models, these indirect effects likely had little impact compared to direct effects. This is consistent with previous studies that found that food availability is the main factor behind the density dependence of growth and body condition (Biro *et al.*, 2003; Amundsen *et al.*, 2007).

One source of uncertainty in our estimates of chub mackerel habitat temperature is our use of the assumption that monthly distribution depth does not change spatiotemporally. In reality, habitat depths can change in response to population fluctuation and environmental variability (Nye *et al.*, 2009). However, we believe that our estimates are reliable. The most frequent habitat of chub mackerel schools and the annual experienced temperatures of chub mackerel in the western North Pacific have been estimated to be about 15–20 °C (Usami, 1973) and 14.4–18.7 °C (Guo *et al.*, 2021), respectively. These temperatures our consistent with our estimates, which provides confidence in our calculations.

The location of the Oyashio Current is also likely to be an important determinant of chub mackerel body condition. Our SEMs showed a positive effect of latitude on body condition and a negative effect of latitude and longitude on habitat temperature (Figure 7 and Table S6). These results are consistent with the strong influence of the Oyashio Current in the northern and eastern waters off northern Japan, and the fact that this current provides an influx of cold water and abundant prey for small pelagic fishes. In addition, our piecewise SEM analysis during the southward migration season showed a strong negative direct effect of SFOI on body condition (Figure 7 and Table S6), meaning that chub mackerel body condition improved with decreasing FOI latitude during the southward migration season. This is a testament to the importance of the FOI as a rich source of nutrients for chub mackerel. The FOI transports a large amount of carbon (in the form of large grazing copepods) from the Oyashio region to the Oyashio– Kuroshio transition region, and this southward transport is estimated to be an important source of organic carbon in this region (Shimizu *et al.*, 2009). It is unclear why the relationships between SFOI, body condition, and temperature during the southward migration season are diametrically opposed to those of the pre-spawning season. One possible factor is the influence of the Kuroshio Current, because almost all samples in the pre-spawning season were caught in waters that are strongly affected by the Kuroshio Current (i.e., <37°N). Future work should examine the effects of the Kuroshio Current on chub mackerel body condition, growth, habitat temperature, and food availability.

Age was also an important endogenous factor for chub mackerel body condition. We identified positive direct and indirect effects of age on body condition during the southward migration and pre-spawning seasons (Figure 7 and Table S6). This suggests that older individuals might distribute in temperatures that are closer to optimum, allowing them to allocate more energy to the accumulation of body energy stores. By contrast, age had a negative direct effect on body condition during the spawning season (Figure 7 and Table S6); this likely reflects the energetic cost of reproduction. Increasing reproductive investment in older, bigger fish (which are more likely to have reached maturity) could explain this decline in body condition.

Previous studies have suggested that the mean growth rate of a given cohort negatively correlates with the population density of the cohort and of the cohort from the preceding year. In chub mackerel, there was negative correlation between size at age 0 (about six months after hatching) and year-class strength (Watanabe and Yatsu, 2004). In Atlantic mackerel, mean length at ages 0–2 negatively correlated with the population density of the cohort and of the cohort from the preceding year (Jansen and Burns, 2015). In our study, the strongest year class of chub mackerel occurred in 2013, and relatively high recruitment continued until 2018 (Table 1). Age-1+ abundance jumped up in 2014 (4.8 times the abundance in 2013) and subsequently remained high, with the 2013 year class accounting for 90% of chub mackerel abundance in 2014 and 57% in 2015 (Table 1). Our results also indicate slower growth of the 2013–2016 year classes after age 0, which is consistent with a previous study (Watanabe and Yatsu, 2004). In addition, growth trajectories and growth-at-age of the 2012 year class indicated slower growth after age 2 (Figure 4 and Table S2). This suggests that growth stagnation of a cohort can be caused by the strength of a subsequent year class, not just by that of the same or previous year.

High growth rates caused early maturation in chub mackerel during the period of declining abundance from 1970 to 2007 (Watanabe and Yatsu, 2006; Watanabe, 2010). Therefore, it is likely that the slow-growing 2013–2016 year classes experienced delayed maturity. Even though high recruitment events occurred several times in the 1990s, recovery of this stock has been prevented due to the overfishing of immature fish (Kawai *et al.*, 2002). To properly maintain and manage this resource, it will be necessary to conserve immature fish and investigate length- and age-at-maturity during the recent phase of increasing stocks.

## Conclusion

Our results show that chub mackerel body condition and growth declined during recent years. These declines were likely caused by the intra- and inter-specific density dependence of these factors. This suggests that intra- and inter-specific competition is a key determinant of chub mackerel body condition and growth variability, and that food availability and carrying capacity for small pelagic fishes in the western North Pacific are approaching their limit. The water temperatures inhabited by chub mackerel were influenced by intra- and inter-specific density dependence, and this likely affected body condition and growth. In other words, our data suggest that the population densities of conspecifics and heterospecifics partly affected chub mackerel body condition and growth through changes in habitat temperature (i.e., indirectly). However, these indirect effects were weaker than the direct effects. This is consistent with previous studies that have identified food availability as the main factor affecting the density dependence of growth and body condition (Biro *et al.*, 2003; Amundsen *et al.*, 2007). Our results will be useful for future stock assessment and management. Further studies of diet and food availability could help clarify the causes of seasonal changes in the intra- and inter-specific density dependence of body condition identified in the present study, and the collection of more data on reproductive traits could help improve our understanding of the plasticity of maturity in chub mackerel.

## Supplementary data

The following supplementary material is available at ICESJMS online.

## Acknowledgements

The authors gratefully acknowledge Masato Kato of the Chiba Prefectural Fisheries Research Center for providing samples and data. We wish to thank Junji Kinoshita for his help in data formatting. The authors also thank English-speaking professional editors from ELSS, Inc., for English proofreading.

## Author contributions

Yasuhiro Kamimura: Conceptualization, Data curation, Methodology, Formal analysis, Investigation, Writing – Original Draft. Makoto Taga: Conceptualization, Investigation, Resources, Writing – Review & Editing. Ryuji Yukami: Data curation, Investigation, Resources, Writing – Review & Editing. Chikako Watanabe: Investigation, Resources, Writing – Review & Editing. Sho Furuichi: Resources and Writing – Review & Editing.

## Data availability

The data that support the findings of this study are available from the corresponding author, YK, upon reasonable request.

## References

Aho, K., Derryberry, D., and Peterson, T. 2014. Model selection for ecologists: the worldviews of AIC and BIC. Ecology, 95 631–636.

Amundsen, P. A., Knudsen, R., and Klemetsen, A. 2007. Intraspecific competition and density dependence of food consumption and growth in Arctic charr. Journal of Animal Ecology, 76 149–158.

Andersen, K. H., Jacobsen, N. S., Jansen, T., and Beyer, J. E. 2017. When in life does density dependence occur in fish populations? Fish and Fisheries, 18 656–667.

Bates, D., Maechler, M., Bolker, B., and Walker, S. 2015. Fitting linear mixed-effects models using lme4. Journal of Statistical Software, 67 1–48.

Biro, P. A., Post, J. R., and Parkinson, E. A. 2003. Density-dependent mortality is mediated by foraging activity for prey fish in whole-lake experiments. Journal of Animal Ecology, 72 546–555.

Blackwell, B. G., Brown, M. L., and Willis, D. W. 2000. Relative weight (Wr) status and current use in fisheries assessment and management. Reviews in Fisheries Science, 8 1–44.

Brosset, P., Ménard, F., Fromentin, J. M., Bonhommeau, S., Ulses, C., Bourdeix, J. H., Bigot, J. L., et al. 2015. Influence of environmental variability and age on the body condition of small pelagic fish in the Gulf of Lions. Marine Ecology Progress Series, 529 219–231.

Brosset, P., Fromentin, J. M., Van Beveren, E., Lloret, J., Marques, V., Basilone, G., Bonanno, A., et al. 2017. Spatio-temporal patterns and environmental controls of small pelagic fish body condition from contrasted Mediterranean areas. Progress in Oceanography, 151 149–162.

Brunel, T., and Dickey-Collas, M. 2010. Effects of temperature and population density on von Bertalanffy growth parameters in Atlantic herring: A macro-ecological analysis. Marine Ecology Progress Series, 405 15–28.

Casini, M., Kornilovs, G., Cardinale, M., Möllmann, C., Grygiel, W., Jonsson, P., Raid, T., et al. 2011. Spatial and temporal density dependence regulates the condition of central Baltic Sea clupeids: Compelling evidence using an extensive international acoustic survey. Population Ecology, 53 511–523.

Casini, M., Käll, F., Hansson, M., Plikshs, M., Baranova, T., Karlsson, O., Lundstrom, K., et al. 2016. Hypoxic areas, density-dependence and food limitation drive the body condition of a heavily exploited marine fish predator. Royal Society Open Science, 3: 160416.

Cormon, X., Ernande, B., Kempf, A., Vermard, Y., and Marchal, P. 2016. North Sea saithe *Pollachius virens* growth in relation to food availability, density dependence and temperature. Marine Ecology Progress Series, 542 141–151.

Dahl, K. A., Edwards, M. A., and Patterson, W. F. 2019. Density-dependent condition and growth of invasive lionfish in the northern Gulf of Mexico. Marine Ecology Progress Series, 623 145–159.

Eero, M., Hjelm, J., Behrens, J., Buchmann, K., Cardinale, M., Casini, M., Gasyukov, P., et al. 2015. Eastern Baltic cod in distress: Biological changes and challenges for stock assessment. ICES Journal of Marine Science, 72 2180–2186.

FAO. 2020. The state of world fisheries and aquaculture 2020. Sustainability in action. Rome. 244 pp.

Froese, R. 2006. Cube law, condition factor and weight-length relationships: History, meta-analysis and recommendations. Journal of Applied Ichthyology, 22 241–253.

Furuichi, S., Yukami, R., Kamimura, Y., Hayashi, A., Isu, S., and Watanabe, R. 2020. Stock assessment and evaluation for the Pacific stock of sardine (fiscal year 2019). *In* Marine Fisheries Stock Assessment and Evaluation for Japanese Waters (Fiscal Year 2019/20).

Guo, C., Ito, S., Yoneda, M., Kitano, H., Kaneko, H., Enomoto, M., Aono, T., et al. 2021. Fish specialize their metabolic performance to maximize bioenergetic efficiency in their local environment: conspecific comparison between two stocks of Pacific chub mackerel (*Scomber japonicus*). Frontiers in Marine Science, 8 1–13.

Higuchi, T., Ito, S., Ishimura, T., Kamimura, Y., Shirai, K., Shindo, H., Nishida, K., et al. 2019. Otolith oxygen isotope analysis and temperature history in early life stages of the chub mackerel *Scomber japonicus* in the Kuroshio-Oyashio transition region. Deep-Sea Research Part II: Topical Studies in Oceanography, 169-170: 104660.

Houde, E. D. 1989. Comparative growth, mortality, and energetics of marine fish larvae: temperature and implied latitudinal effects. Fishery Bulletin, 87: 471–495.

Hunter, J. R., and Kimbrell, C. A. 1980. Early life history of Pacific mackerel, Scomber 592 japonicus. Fisheries Bulletin, 78: 89–100.

Itoh, S., Yasuda, I., Saito, H., Tsuda, A., and Komatsu, K. 2015. Mixed layer depth and chlorophyll a: Profiling float observations in the Kuroshio-Oyashio Extension region. Journal of Marine Systems, 151 1–14.

James, G., Witten, D., Hastie, T., and Tibshirani, R. 2013. An Introduction to Statistical Learning: with Applications in R. Springer-Verlag New York. 426 pp.

Jansen, T., and Burns, F. 2015. Density dependent growth changes through juvenile and early adult life of North East Atlantic Mackerel (*Scomber scombrus*). Fisheries Research, 169 37–44.

Kamimura, Y., Takahashi, M., Yamashita, N., Watanabe, C., and Kawabata, A. 2015. Larval and juvenile growth of chub mackerel *Scomber japonicus* in relation to recruitment in the western North Pacific. Fisheries Science, 81 505–513.

Kanamori, Y., Takasuka, A., Nishijima, S., and Okamura, H. 2019. Climate change shifts the spawning ground northward and extends the spawning period of chub mackerel in the western North Pacific. Marine Ecology Progress Series, 624 155–166.

Kaneko, H., Okunishi, T., Seto, T., Kuroda, H., Itoh, S., Kouketsu, S., and Hasegawa, D. 2019. Dual effects of reversed winter-spring temperatures on year-to-year variation in the recruitment of chub mackerel (*Scomber japonicus*). Fisheries Oceanography, 28 212–227.

Kawai, H., Yatsu, A., Watanabe, C., Mitani, T., Katsukawa, T., and Matsuda, H. 2002. Recovery policy for chub mackerel stock using recruitment-per-spawning. Fisheries Science, 68 963–971.

Kawasaki, T. 1984. Food habits of the far eastern sardine and their implication in the fluctuation pattern of the sardine stocks. Bulletin of the Japanese Society of Scientific Fisheries, 50: 1657–1663 (in Japanese with English abstract).

Kawashima, T., Ishi, M., and Katayama, S. 2017. Age determination of chub mackerel *Scomberjaponicus* using otolith transverse section. Journal of Fisheries Technology, 9: 45–51 (in Japanese with English abstract).

Kishida, T., and Matsuda, H. 1993. Statistical analyses of intra J and interspecific density effects on recruitment of chub mackerel and sardine in Japan. Fisheries Oceanography, 2 278–287.

Kobari, T., Nagaki, T., and Takahashi, K. 2004. Seasonal changes in abundance and development of *Calanuspacificus* (Crustacea: Copepoda) in the Oyashio-Kuroshio Mixed Region. Marine Biology, 144 713–721.

Kobari, T., Moku, M., and Takahashi, K. 2008. Seasonal appearance of expatriated boreal copepods in the Oyashio-Kuroshio mixed region. ICES Journal of Marine Science, 65 469–476.

Kondo, K., and Kuroda, K. 1966. Growth of the Japanese mackerel-I. Comparison of resting zones on different organs. Bulletin of Tokai Regional Fisheries Research Laboratory, 45: 31–60 (in Japanese with English abstract).

Kuroda, H., Setou, T., Kakehi, S., Ito, S., Taneda, T., Azumaya, T., Inagake, D., et al. 2017. Recent advances in Japanese fisheries science in the Kuroshio-Oyashio region through development of the FRA-ROMS ocean forecast system: Overview of the reproducibility of reanalysis products. Open Journal of Marine Science, 7 62–90.

Le cren, E. D. 1951. The length-weight relationship and seasonal cycle in gonad weight and condition in the perch (*Perca fluviatilis*). The Journal of Animal Ecology, 20 201–219.

Lefcheck, J. S. 2016. PIECEWISESEM: Piecewise structural equation modelling in R for ecology, evolution, and systematics. Methods in Ecology and Evolution, 7 573–579.

Lloret, J., Gil de Sola, L., Souplet, A., and Galzin, R. 2002. Effects of large-scale habitat variability on condition of demersal exploited fish in the north-western Mediterranean. ICES Journal of Marine Science, 59 1215–1227.

Lloret, J., Shulman, G., and Love, R. M. 2014. Condition and Health Indicators of Exploited Marine Fishes. Wiley-Blackwell, West Sussex, UK. 262 pp.

Lorenzen, K., and Enberg, K. 2002. Density-dependent growth as a key mechanism in the regulation of fish populations: Evidence from among-population comparisons. Proceedings of the Royal Society B: Biological Sciences, 269 49–54.

Lorenzen, K. 2008. Fish population regulation beyond ‘stock and recruitment’: The role of density-dependent growth in the recruited stock. Bulletin of Marine Science, 83 181–196.

Murakami, M. 1994. On long-term variations in hydrographic conditions in the Tohoku area. Bulletin of Tohoku Regional Fisheries Research Institute, 56: 47–56 (in Japanese with English abstract).

Nakai, Z. 1962. Studies relevant sardine to mechanisms underlying the fluctuation in the catch of the Japanese sardine *Sardinops melanosticta* (TEMMINCK & SCHLEGEL). Japanese Journal of Ichthyology, 9 1–115.

Nakamura, M., Yoneda, M., Ishimura, T., Shirai, K., Tamamura, M., and Nishida, K. 2020. Temperature dependency equation for chub mackerel (*Scomberjaponicus*) identified by a laboratory rearing experiment and microscale analysis. Marine and Freshwater Research, 71 1384–1389.

Nakatsuka, S., Kawabata, A., Takasuka, A., Kubota, H., Okamura, H., and Oozeki, Y. 2010. Estimating gastric evacuation rate and daily ration of chub mackerel and spotted mackerel in the Kuroshio-Oyashio transition and Oyashio regions. Bulletin of Japanese Society of Fisheries Oceanography, 74: 105–117 (in Japanese with English abstract).

Nye, J. A., Link, J. S., Hare, J. A., and Overholtz, W. J. 2009. Changing spatial distribution of fish stocks in relation to climate and population size on the Northeast United States continental shelf. Marine Ecology Progress Series, 393 111–129.

Ogle, D. H., Wheeler, P., and Dinno, A. 2020. FSA: Fisheries Stock Analysis. https://github.com/droglenc/FSA.

Olafsdottir, A. H., Slotte, A., Jacobsen, J. A., Oskarsson, G. J., Utne, K. R., and Nottestad, L. 2016. Changes in weight-at-length and size-at-age of mature Northeast Atlantic mackerel (*Scomber scombrus*) from 1984 to 2013: effects of mackerel stock size and herring (*Clupea harengus*) stock size. ICES Journal of Marine Science, 73 1255–1265.

Parrish, R. H., and Mallicoate, D. L. 1995. Variation in the condition factors of California pelagic fishes and associated environmental factors. Fisheries Oceanography, 4 171–190.

Qiu, B. 2001. Kuroshio and Oyashio Currents. Encyclopedia of Ocean Sciences: 1413-1425.

R Core Team. 2020. R: A Language and Environment for Statistical Computing. R Foundation for Statistical Computing, Vienna, Austria. https://www.r-project.org/.

Rose, K. A., Cowan, J. H., Winemiller, K. O., Myers, R. A., and Hilborn, R. 2001. Compensatory density dependence in fish populations: Importance, controversy, understanding and prognosis. Fish and Fisheries, 2 293–327.

Rueda, L., Assutí, E. M., Alvarez-Berastegu, D., and Hidalgo, M. 2015. Effect of intra-specific competition, surface chlorophyll and fishing on spatial variation of gadoid’s body condition. Ecosphere, 6 1–17.

Sassa, C., and Tsukamoto, Y. 2010. Distribution and growth of *Scomber japonicus* and *S. australasicus* larvae in the southern East China Sea in response to oceanographic conditions. Marine Ecology Progress Series, 419 185–199.

Sato, Y. 1974. Environmental factors affecting to the distribution of the mackerel shoals in the doto and the sanriku fishing grounds, north eastern sea of Japan (1) surface water temperatures at the fishing spots. Bulletin of Tohoku Regional Fisheries Research Laboratory, 34: 17–30 (in Japanese with English abstract).

Sato, Y., Iizuka, K., and Kubota, S. 1975. Some features of the inter-specific relations between mackerel and other fishes distributed in the fishing grounds in the north-eastern sea of Japan. Bulletin of Tohoku Regional Fisheries Research Laboratory, 35: 1–13 (in Japanese with English abstract).

Shimizu, Y., Takahashi, K., Ito, S. I., Kakehi, S., Tatebe, H., Yasuda, I., Kusaka, A., et al. 2009. Transport of subarctic large copepods from the Oyashio area to the mixed water region by the coastal Oyashio intrusion. Fisheries Oceanography, 18 312–327.

Shipley, B. 2000. A new inferential test for path models based on directed acyclic graphs. Structural Equation Modeling, 7 206–218.

Shipley, B. 2009. Confirmatory path analysis in a generalized multilevel context. Ecology, 90 363–368.

Swain, D. P., and Kramer, D. L. 1995. Annual variation in temperature selection by Atlantic cod *Gadus morhua* in the southern Gulf of St. Lawrence, Canada, and its relation to population size. Marine Ecology Progress Series, 116 11–24.

Taga, M., and Yamashita, Y. 2018. Evaluation of incidental ingestion by stomach contents analysis of chub mackerel *Scomber japonicus* caught by purse seine. Bulletin of Japanese Society of Fisheries Oceanography, 82: 121–126 (in Japanese with English abstract).

Taga, M., Kamimura, Y., and Yamashita, Y. 2019. Effects of water temperature and prey density on recent growth of chub mackerel *Scomber japonicus* larvae and juveniles along the Pacific coast of Boso-Kashimanada. Fisheries Science, 85 931–942.

Takahashi, M., Yoneda, M., Kitano, H., Kawabata, A., and Saito, M. 2014. Growth of juvenile chub mackerel *Scomber japonicus* in the western North Pacific Ocean: with application and validation of otolith daily increment formation. Fisheries Science, 80 293–300.

Takasuka, A., Yoneda, M., and Oozeki, Y. 2020. Density-dependent egg production in chub mackerel in the Kuroshio Current system. Fisheries Oceanography: 1-13.

Usami, S. 1973. Ecological studies of life pattern of the Japanese mackerel, *Scomber japonicus* HOUTTUYN on the adult of the Pacific subpopulation. Bulletin of Tokai Regional Fisheries Research Laboratory, 76: 71–178 (in Japanese with English abstract).

Van Gemert,R., and Andersen, K. H. 2018. Implications of late-in-life density-dependent growth for fishery size-at-entry leading to maximum sustainable yield. ICES Journal of Marine Science, 75 1296–1305.

Watanabe, C., and Nishida, H. 2002. Development of assessment techniques for pelagic fish stocks: applications of daily egg production method and pelagic trawl in the Northwestern Pacific Ocean. Fisheries Science: 97-100.

Watanabe, C., and Yatsu, A. 2004. Effects of density-dependence and sea surface temperature on interannual variation in length-at-age of chub mackerel (*Scomber japonicus*) in the Kuroshio-Oyashio area during 1970-1997. Fisheries Bulletin, 102:196–206.

Watanabe, C., and Yatsu, A. 2006. Long-term changes in maturity at age of chub mackerel (*Scomber japonicus*) in relation to population declines in the waters off northeastern Japan. Fisheries Research, 78 323–332.

Watanabe, C. 2010. Changes in the reproductive traits of the Pacific stock of chub mackerel *Scomber japonicus* and their effects on the population dynamics. Bulletin of the Japanese Society of Fisheries Oceanography, 74: 46–50 (in Japanese with English abstract).

Watanabe, T. 1970. Morphology and ecology of early stages of life in Japanese common mackerel, *Scomber japonicus* HOUTTYN, with special reference to fluctuation of population. Bulletin of Tokai Regional Fisheries Research Laboratory, 62: 1–283 (in Japanese with English abstract).

Yamamoto, M., and Katayama, S. 2012. Interspecific comparisons of feeding habit between Japanese anchovy *Engraulis japonicus* and Japanese sardine *Sardinops melanostictus* in eastern Hiuchi-nada, central Seto Inland Sea, Japan, in 1995. Bull. Jpn. Soc. Fish. Oceanogr., 76: 66–76 (in Japanese with English abstract).

Yamashita, H. 1955. The feeding habit of sardine, *sardinia melanosticta,* in the waters adjacent to Kyusyu, with reference to its growth. Bulletin of the Japanese Society of Scientific Fisheries, 21: 471–475 (in Japanese with English abstract).

Yasuda, I. 2003. Hydrographic structure and variability in the Kuroshio-Oyashio transition area. Journal of Oceanography, 59 389–402.

Yatsu, A., Watanabe, T., Ishida, M., Sugisaki, H., and Jacobson, L. D. 2005. Environmental effects on recruitment and productivity of Japanese sardine *Sardinops melanostictus* and chub mackerel Scomber japonicus with recommendations for management. Fisheries Oceanography, 14 263–278.

Yatsu, A., Chiba, S., Yamanaka, Y., Ito, S. I., Shimizu, Y., Kaeriyama, M., and Watanabe, Y. 2013. Climate forcing and the Kuroshio/Oyashio ecosystem. ICES Journal of Marine Science, 70 922–933.

Yatsu, A., Takahashi, K., Watanabe, K., and Honda, O. 2019. Seasonal and interannual variability in crude fat content and condition factor of chub mackerel Scomber japonicus captured from waters off northeastern Japan during 2012 J2017. Bull. Jpn. Soc. Fish. Oceanogr., 83: 19–27 (in Japanese with English abstract).

Yatsu, A. 2019. Review of population dynamics and management of small pelagic fishes around the Japanese Archipelago. Fisheries Science, 85 611–639.

Yukami, R., Nishijima, S., Isu, S., Kamimura, Y., Furuichi, S., and Watanabe, R. 2020a. Stock assessment and evaluation for Pacific stock of chub mackerel (fisical year 2019). *In* Marine fisheries stock assessment and evaluation for Japanese waters (fiscal year 2019/2020. Fisheries Agency and Fisheries Research and Education Agency of Japan, Tokyo.

Yukami, R., Isu, S., Kamimura, Y., Furuichi, S., Watanabe, R., and Kanamori, Y. 2020b. Stock assessment and evaluation for Pacific stock of spotted mackerel (fisical year 2019). *In* Marine fisheries stock assessment and evaluation for Japanese waters (fiscal year 2019/2020. Fisheries Agency and Fisheries Research and Education Agency of Japan.

Zimmermann, F., Ricard, D., and Heino, M. 2018. Density regulation in Northeast Atlantic fish populations: Density dependence is stronger in recruitment than in somatic growth. Journal of Animal Ecology, 87 672–681.

