## Supplementary material for "Intra- and inter-specific density dependence of body condition, growth, and habitat temperature in chub mackerel (*Scomber japonicus*)"

Table S1. Estimated parameters of von Bertalanffy growth formulae by sex and year class. Comparisons among sexes were made by using *F* tests.

| Year class | Sex | *n* | $L_{inf}$ | *K* | $t_0$ | *F* value | *p*-value |
|---|---|---|---|---|---|---|---|
| 2006 | Male | 219 | 495.6 | 0.17 | −3.39 | 0.08 | 0.97 |
| | Female | 236 | 500.5 | 0.17 | −3.27 | | |
| 2007 | Male | 575 | 398.8 | 0.49 | −1.31 | 1.37 | 0.25 |
| | Female | 609 | 405.2 | 0.48 | −1.25 | | |
| 2008 | Male | 273 | 395.5 | 0.51 | −1.31 | 3.79 | 0.01 |
| | Female | 283 | 401.2 | 0.47 | −1.58 | | |
| 2009 | Male | 408 | 402.8 | 0.43 | −1.60 | 3.96 | 0.01 |
| | Female | 531 | 405.3 | 0.42 | −1.70 | | |
| 2010 | Male | 453 | 393.3 | 0.51 | −1.54 | 2.53 | 0.06 |
| | Female | 526 | 399.7 | 0.49 | −1.45 | | |
| 2011 | Male | 286 | 387.6 | 0.41 | −2.53 | 1.71 | 0.16 |
| | Female | 352 | 387.5 | 0.47 | −2.05 | | |
| 2012 | Male | 638 | 394.8 | 0.22 | −4.91 | 0.81 | 0.49 |
| | Female | 757 | 398.2 | 0.20 | −5.19 | | |
| 2013 | Male | 1 799 | 362.5 | 0.27 | −3.06 | 2.87 | 0.04 |
| | Female | 2 066 | 372.1 | 0.26 | −3.04 | | |
| 2014 | Male | 602 | 354.9 | 0.31 | −2.91 | 1.60 | 0.19 |
| | Female | 746 | 337.6 | 0.45 | −1.95 | | |
| 2015 | Male | 217 | 388.1 | 0.29 | −1.89 | 0.13 | 0.94 |
| | Female | 287 | 415.1 | 0.23 | −2.45 | | |
| 2016 | Male | 183 | 368.3 | 0.35 | −1.77 | 1.16 | 0.32 |
| | Female | 221 | 327.9 | 0.51 | −1.43 | | |

Table S2. Estimated fork length (FL) at ages 1−6 and annual growth rate by length ($G_L$) at ages 1−5 by year class, calculated by using the von Bertalanffy growth formula.

| Estimate | Year | Age | | | | | |
|---|---|---|---|---|---|---|---|
| | class | 1 | 2 | 3 | 4 | 5 | 6 |
| FL | 2006 | 253.5 | 296.8 | 330.1 | 355.7 | 375.3 | 390.4 |
| | 2007 | 268.7 | 319.9 | 351.2 | 370.3 | 382.0 | 389.1 |
| | 2008 | 275.6 | 323.0 | 351.7 | 369.2 | 379.7 | 386.1 |
| | 2009 | 273.0 | 318.2 | 347.4 | 366.3 | 378.4 | 386.3 |
| | 2010 | 281.4 | 325.1 | 352.2 | 369.0 | 379.5 | 385.9 |
| | 2011 | 294.4 | 324.0 | 344.7 | 359.3 | 369.5 | 376.7 |
| | 2012 | 264.9 | 305.4 | 328.7 | 342.1 | 349.8 | 354.2 |
| | 2013 | 235.3 | 268.3 | 292.3 | 309.8 | 322.6 | 331.9 |
| | 2014 | 245.4 | 276.7 | 298.5 | 313.6 | 324.2 | - |
| | 2015 | 224.7 | 264.3 | 294.9 | 318.6 | - | - |
| | 2016 | 224.2 | 267.7 | 294.8 | - | - | - |
| $G_L$ | 2006 | 43.3 | 33.3 | 25.6 | 19.6 | 15.1 | - |
| | 2007 | 51.2 | 31.3 | 19.1 | 11.7 | 7.2 | - |
| | 2008 | 47.4 | 28.7 | 17.4 | 10.6 | 6.4 | - |
| | 2009 | 45.2 | 29.2 | 18.9 | 12.2 | 7.9 | - |
| | 2010 | 43.7 | 27.1 | 16.8 | 10.4 | 6.5 | - |
| | 2011 | 29.6 | 20.8 | 14.6 | 10.2 | 7.2 | - |
| | 2012 | 40.5 | 23.3 | 13.4 | 7.7 | 4.4 | - |
| | 2013 | 33.0 | 24.0 | 17.5 | 12.8 | 9.3 | - |
| | 2014 | 31.3 | 21.8 | 15.2 | 10.6 | - | - |
| | 2015 | 39.5 | 30.6 | 23.7 | - | - | - |
| | 2016 | 43.5 | 27.1 | - | - | - | - |

Table S3. The number of chub mackerel and the ranges of variables used for piecewise structural equation models (SEMs) by quarter and year. Abbreviations are as follows: $K_n$, relative condition factor; TC, habitat temperature; $N_m$, chub mackerel abundance; $N_s$, Japanese sardine abundance; $A$, age; SFOI, the southernmost position of the first Oyashio intrusion; Lat, catch latitude; Lon, catch longitude.

| Quarter | Year | $n$ | $K_n$ | TC | $N_m$ | $N_s$ | $A$ | SFOI | Lat | Lon |
|---|---|---|---|---|---|---|---|---|---|---|
| Q1 | 2007 | 8 | 0.91–1.12 | 17.4–17.6 | 0.85 | 1.09 | 1 | 38.8–39.1 | 34.9–35.9 | 140–140.9 |
| Q1 | 2008 | 173 | 0.9–1.26 | 12.7–21.8 | 1.08 | 0.52 | 1–2 | 38.9–40 | 34.5–36.1 | 139.3–141.1 |
| Q1 | 2009 | 174 | 0.86–1.23 | 13.9–23 | 0.71 | 1.41 | 1–3 | 39.8–40.1 | 34.1–36.8 | 139.4–141.6 |
| Q1 | 2010 | 250 | 0.82–1.24 | 13.4–22.8 | 1.34 | 1.17 | 1–4 | 39.5–39.6 | 34.5–36 | 139.3–141.3 |
| Q1 | 2011 | 61 | 0.82–1.1 | 17.4–20.9 | 1.46 | 5.25 | 2–5 | 40–40.3 | 34.5–35.8 | 139.4–141.3 |
| Q1 | 2012 | 260 | 0.77–1.18 | 17.2–22.1 | 1.57 | 4.90 | 1–6 | 39.8–40.5 | 34.2–36.1 | 138.5–141.5 |
| Q1 | 2013 | 150 | 0.8–1.04 | 19.1–21.3 | 2.88 | 6.27 | 2–5 | 38.4–39 | 34.3–34.5 | 139–139.4 |
| Q1 | 2014 | 139 | 0.8–1.17 | 14.8–21.1 | 13.83 | 7.72 | 1–7 | 38.3–39.3 | 35.7–36.7 | 140.9–141.6 |
| Q1 | 2015 | 289 | 0.77–1.12 | 11.3–21.3 | 12.29 | 15.92 | 1–7 | 39.1–39.9 | 33.5–36.4 | 139.5–141.7 |
| Q1 | 2016 | 329 | 0.78–1.37 | 11.3–21.8 | 10.27 | 30.96 | 1–7 | 40–41.2 | 33.8–36.7 | 138.5–141.5 |
| Q1 | 2017 | 362 | 0.75–1.24 | 13.9–21.1 | 14.20 | 31.40 | 1–9 | 38.2–39.1 | 33.9–38.2 | 138.6–141.6 |
| Q1 | 2018 | 276 | 0.71–1.09 | 13.3–21.1 | 14.12 | 31.36 | 2–10 | 39.7–40.2 | 34.1–38.5 | 138.7–141.7 |
| Q2 | 2007 | 1 | 0.96 | 21.8 | 0.85 | 1.09 | 1 | 39.6 | 35.6 | 140.9 |
| Q2 | 2008 | 28 | 0.94–1.22 | 18.2–23.1 | 1.08 | 0.52 | 1–2 | 40.1–40.6 | 35.8–38.9 | 141.3–141.9 |
| Q2 | 2009 | 157 | 0.92–1.33 | 17.2–19.8 | 0.71 | 1.41 | 1–3 | 38.5–40 | 37.8–40.8 | 141.6–142 |
| Q2 | 2010 | 231 | 0.85–1.35 | 13.4–25.2 | 1.34 | 1.17 | 1–4 | 39.3–39.7 | 35.8–41.9 | 141.2–143.5 |

| | | | | | | | | | | |
|---|---|---|---|---|---|---|---|---|---|---|
| Q2 | 2011 | 84 | 0.86–1.34 | 18.8–21.3 | 1.46 | 5.25 | 1–5 | 39.8–41.2 | 35.8–40.3 | 141.1–142.1 |
| Q2 | 2014 | 160 | 0.81–1.21 | 14.1–21.6 | 13.83 | 7.72 | 1–3 | 39.4–40.3 | 39.2–42.8 | 141.7–144.7 |
| Q2 | 2015 | 108 | 0.82–1.25 | 16.5–21 | 12.29 | 15.92 | 1–5 | 40–40.1 | 35.8–40.9 | 141–142 |
| Q2 | 2016 | 51 | 0.91–1.08 | 17.9–22.7 | 10.27 | 30.96 | 1–3 | 40.1–40.5 | 35.8–40.8 | 141–141.8 |
| Q2 | 2017 | 116 | 0.81–1.21 | 9.7–20.1 | 14.20 | 31.40 | 2–5 | 39.5 | 40.7–42.8 | 141.7–144.6 |
| Q2 | 2018 | 131 | 0.85–1.32 | 18.5–21.5 | 14.12 | 31.36 | 2–6 | 39.7–39.8 | 40.6–40.8 | 141.6–145 |
| Q3 | 2007 | 5 | 0.96–1.09 | 18.7–22.2 | 0.85 | 1.09 | 1 | 41.8 | 35.6–36.7 | 141–141.2 |
| Q3 | 2008 | 160 | 0.83–1.32 | 14.3–21.6 | 1.08 | 0.52 | 1–2 | 40.9–41.3 | 34.9–40.6 | 139.9–142 |
| Q3 | 2009 | 81 | 0.9–1.37 | 15.7–19 | 0.71 | 1.41 | 1–3 | 39.9–40.7 | 36.7–40.6 | 140.9–142.2 |
| Q3 | 2010 | 155 | 0.82–1.32 | 18.7–20.6 | 1.34 | 1.17 | 1–4 | 40.2–40.7 | 36.8–40.4 | 141.2–142.2 |
| Q3 | 2011 | 53 | 0.95–1.32 | 18.6–21.2 | 1.46 | 5.25 | 1–5 | 39.7–40.8 | 35.5–40.4 | 141.2–142.1 |
| Q3 | 2012 | 158 | 0.89–1.29 | 18.2–23.7 | 1.57 | 4.90 | 1–6 | 40.7–41.3 | 35.8–38.2 | 140.9–141.7 |
| Q3 | 2013 | 54 | 0.92–1.34 | 15.3–18.3 | 2.88 | 6.27 | 1–5 | 41.7 | 35.8–36.8 | 141.1–142.4 |
| Q3 | 2014 | 256 | 0.89–1.25 | 13.5–19.3 | 13.83 | 7.72 | 1–5 | 40.4–41.2 | 36.3–40.6 | 141–142.4 |
| Q3 | 2015 | 465 | 0.85–1.24 | 14.9–18.6 | 12.29 | 15.92 | 1–6 | 40.1–42.4 | 35.8–40.1 | 141.1–142.4 |
| Q3 | 2016 | 436 | 0.8–1.3 | 12.5–22.9 | 10.27 | 30.96 | 1–8 | 41–41.7 | 35.4–42.8 | 140.9–145 |
| Q3 | 2017 | 264 | 0.82–1.38 | 14.7–21.6 | 14.20 | 31.40 | 1–7 | 40.3–41.4 | 35.6–40.2 | 141–142.2 |
| Q3 | 2018 | 282 | 0.8–1.3 | 16.5–19.1 | 14.12 | 31.36 | 2–8 | 39.9–40 | 36.8–40.6 | 141.2–142.2 |
| Q4 | 2008 | 28 | 0.82–1.2 | 14.7–17.6 | 1.08 | 0.52 | 1 | 39.1–40.3 | 34.3–37 | 139–141.3 |
| Q4 | 2009 | 137 | 0.85–1.24 | 10.1–18 | 0.71 | 1.41 | 1–2 | 39.3–39.8 | 34.3–37 | 139–141.3 |
| Q4 | 2010 | 158 | 0.84–1.16 | 13.6–17.9 | 1.34 | 1.17 | 1–3 | 37.7–39.7 | 34.1–35.8 | 139–141.2 |
| Q4 | 2011 | 111 | 0.85–1.29 | 11.3–17.9 | 1.46 | 5.25 | 1–4 | 37.6–40.6 | 34.3–35.9 | 139–141.2 |
| Q4 | 2012 | 162 | 0.82–1.27 | 13.8–19.1 | 1.57 | 4.90 | 1–4 | 38.8–39.9 | 33.9–35.9 | 138.8–141.3 |

| | | | | | | | | | | |
|---|---|---|---|---|---|---|---|---|---|---|
| Q4 | 2013 | 63 | 0.88–1.24 | 13.6–17.1 | 2.88 | 6.27 | 1–3 | 39.4–40.3 | 35.8–36.6 | 141–141.1 |
| Q4 | 2014 | 103 | 0.82–1.25 | 12.2–16.3 | 13.83 | 7.72 | 1–7 | 38.9–40.6 | 35.8–36.8 | 140.9–141.2 |
| Q4 | 2015 | 283 | 0.77–1.17 | 9.9–17.1 | 12.29 | 15.92 | 1–5 | 39–40.2 | 34.1–36.4 | 139.4–141.2 |
| Q4 | 2016 | 649 | 0.77–1.15 | 9.4–19.2 | 10.27 | 30.96 | 1–8 | 40–42 | 33.8–36.8 | 138.3–141.2 |
| Q4 | 2017 | 505 | 0.73–1.22 | 7.6–18.8 | 14.20 | 31.40 | 1–8 | 37.4–41.4 | 33.7–36.7 | 138.3–141.2 |
| Q4 | 2018 | 323 | 0.76–1.21 | 10.3–16.2 | 14.12 | 31.36 | 1–5 | 39.6–40.7 | 34–36.6 | 138.5–141.2 |

Table S4. The number of chub mackerel and the means and standard deviations of variables used for piecewise structural equation models (SEMs) by quarter and year. Abbreviations are as follows: $K_n$, relative condition factor; TC, habitat temperature; SFOI, the southernmost position of the first Oyashio intrusion; Lat, catch latitude; Lon, catch longitude.

| Quarter | Year | *n* | $K_n$ | TC | SFOI | Lat | Lon |
|---|---|---|---|---|---|---|---|
| Q1 | 2007 | 8 | 1.01 ± 0.06 | 17.4 ± 0.1 | 39 ± 0.1 | 35.1 ± 0.5 | 140.2 ± 0.4 |
| Q1 | 2008 | 173 | 1.05 ± 0.07 | 19.7 ± 2.4 | 39.6 ± 0.4 | 35.2 ± 0.6 | 140.4 ± 0.8 |
| Q1 | 2009 | 174 | 1.01 ± 0.07 | 18.4 ± 2.7 | 40 ± 0.1 | 35.5 ± 0.9 | 140.6 ± 0.9 |
| Q1 | 2010 | 250 | 0.97 ± 0.07 | 19.1 ± 2.5 | 39.5 ± 0.1 | 35.3 ± 0.7 | 140.4 ± 0.8 |
| Q1 | 2011 | 61 | 0.94 ± 0.06 | 20.1 ± 1.3 | 40.1 ± 0.1 | 35 ± 0.6 | 140.2 ± 0.9 |
| Q1 | 2012 | 260 | 0.95 ± 0.06 | 20.2 ± 1.6 | 40.2 ± 0.2 | 35.2 ± 0.7 | 140.2 ± 1.1 |
| Q1 | 2013 | 150 | 0.91 ± 0.04 | 20.3 ± 0.8 | 38.8 ± 0.3 | 34.5 ± 0.1 | 139.3 ± 0.1 |
| Q1 | 2014 | 139 | 0.97 ± 0.07 | 17.1 ± 1.7 | 38.7 ± 0.4 | 36.3 ± 0.3 | 141.2 ± 0.2 |
| Q1 | 2015 | 289 | 0.91 ± 0.06 | 16.3 ± 3.3 | 39.3 ± 0.3 | 35.5 ± 1 | 140.7 ± 0.7 |
| Q1 | 2016 | 329 | 0.92 ± 0.07 | 16.2 ± 2.9 | 40.4 ± 0.4 | 35.7 ± 0.8 | 140.8 ± 0.9 |
| Q1 | 2017 | 362 | 0.91 ± 0.06 | 16.1 ± 1.7 | 38.9 ± 0.3 | 36 ± 1 | 140.7 ± 0.8 |
| Q1 | 2018 | 276 | 0.92 ± 0.06 | 19.1 ± 2 | 40.1 ± 0.2 | 34.9 ± 0.8 | 140 ± 0.9 |
| Q2 | 2007 | 1 | 0.96 | 21.8 | 39.6 | 35.6 | 140.9 |
| Q2 | 2008 | 28 | 1.06 ± 0.06 | 22.1 ± 1.9 | 40.1 ± 0.1 | 36.3 ± 1.1 | 141.3 ± 0.2 |
| Q2 | 2009 | 157 | 1.11 ± 0.07 | 18.2 ± 0.7 | 39.4 ± 0.7 | 40.4 ± 0.8 | 141.8 ± 0.1 |
| Q2 | 2010 | 231 | 1.13 ± 0.11 | 18.2 ± 3.1 | 39.6 ± 0.2 | 40.4 ± 1.5 | 142.2 ± 0.7 |

| | | | | | | | |
|---|---|---|---|---|---|---|---|
| Q2 | 2011 | 84 | 1.07 ± 0.1 | 19.3 ± 0.9 | 40 ± 0.5 | 39.6 ± 1.6 | 141.9 ± 0.3 |
| Q2 | 2014 | 160 | 1.06 ± 0.06 | 19.7 ± 1.8 | 39.7 ± 0.3 | 40.8 ± 0.2 | 141.8 ± 0.3 |
| Q2 | 2015 | 108 | 0.99 ± 0.06 | 18.7 ± 1.4 | 40 ± 0 | 39 ± 2.3 | 141.6 ± 0.4 |
| Q2 | 2016 | 51 | 0.99 ± 0.04 | 18.9 ± 1.3 | 40.3 ± 0.2 | 40.3 ± 1.5 | 141.6 ± 0.2 |
| Q2 | 2017 | 116 | 1.02 ± 0.07 | 15.3 ± 4.7 | 39.5 ± 0 | 41.6 ± 1 | 142.9 ± 1.4 |
| Q2 | 2018 | 131 | 1.04 ± 0.07 | 20.1 ± 1.1 | 39.7 ± 0 | 40.8 ± 0.1 | 142.7 ± 1.6 |
| Q3 | 2007 | 5 | 1.02 ± 0.05 | 20.8 ± 1.9 | 41.8 ± 0 | 36 ± 0.6 | 141.1 ± 0.1 |
| Q3 | 2008 | 160 | 1.05 ± 0.1 | 18 ± 2.4 | 41.1 ± 0.2 | 37.1 ± 1.5 | 141.3 ± 0.6 |
| Q3 | 2009 | 81 | 1.14 ± 0.11 | 17.3 ± 1.3 | 40.2 ± 0.4 | 38.5 ± 1.2 | 141.6 ± 0.3 |
| Q3 | 2010 | 155 | 1.11 ± 0.09 | 19.7 ± 0.5 | 40.3 ± 0.2 | 39.4 ± 1.4 | 141.9 ± 0.3 |
| Q3 | 2011 | 53 | 1.13 ± 0.09 | 20.1 ± 1 | 40 ± 0.5 | 38.7 ± 2.3 | 141.7 ± 0.4 |
| Q3 | 2012 | 158 | 1.03 ± 0.08 | 20.3 ± 1.5 | 40.9 ± 0.2 | 36.5 ± 1 | 141.3 ± 0.3 |
| Q3 | 2013 | 54 | 1.09 ± 0.09 | 17 ± 1.3 | 41.7 ± 0 | 36.7 ± 0.2 | 141.8 ± 0.6 |
| Q3 | 2014 | 256 | 1.03 ± 0.07 | 16.5 ± 1.9 | 40.8 ± 0.3 | 38.8 ± 1.2 | 142 ± 0.3 |
| Q3 | 2015 | 465 | 0.99 ± 0.06 | 16.5 ± 1.1 | 41.7 ± 0.8 | 38.6 ± 1.3 | 141.8 ± 0.5 |
| Q3 | 2016 | 436 | 1.05 ± 0.08 | 16.9 ± 2.2 | 41.4 ± 0.3 | 38 ± 2 | 141.8 ± 1.1 |
| Q3 | 2017 | 264 | 1.09 ± 0.09 | 16.7 ± 1.6 | 41 ± 0.4 | 38.5 ± 1.5 | 141.8 ± 0.4 |
| Q3 | 2018 | 282 | 1.07 ± 0.09 | 17.6 ± 0.8 | 39.9 ± 0.1 | 38.8 ± 1.4 | 141.8 ± 0.3 |
| Q4 | 2008 | 28 | 0.98 ± 0.08 | 16 ± 0.7 | 40 ± 0.4 | 35.6 ± 1 | 140.4 ± 0.8 |
| Q4 | 2009 | 137 | 1.03 ± 0.09 | 14.2 ± 1.9 | 39.6 ± 0.2 | 35.4 ± 1 | 140.2 ± 1 |
| Q4 | 2010 | 158 | 0.99 ± 0.07 | 15.4 ± 1.2 | 38.9 ± 0.8 | 35.2 ± 0.7 | 140.6 ± 0.9 |
| Q4 | 2011 | 111 | 1.02 ± 0.09 | 16 ± 1.8 | 38.8 ± 1.4 | 34.9 ± 0.7 | 140 ± 1 |
| Q4 | 2012 | 162 | 1.03 ± 0.08 | 17.1 ± 1.3 | 39.4 ± 0.4 | 34.6 ± 0.6 | 139.4 ± 0.9 |

| | | | | | | | |
|---|---|---|---|---|---|---|---|
| Q4 | 2013 | 63 | 1.02 ± 0.09 | 15.6 ± 1.1 | 40.1 ± 0.4 | 36.1 ± 0.3 | 141 ± 0 |
| Q4 | 2014 | 103 | 0.98 ± 0.07 | 14.2 ± 1.4 | 39.9 ± 0.6 | 36.2 ± 0.4 | 141.1 ± 0.1 |
| Q4 | 2015 | 283 | 0.95 ± 0.08 | 14.1 ± 2.5 | 39.6 ± 0.5 | 35.5 ± 0.8 | 140.6 ± 0.7 |
| Q4 | 2016 | 649 | 0.92 ± 0.06 | 14 ± 2.5 | 40.9 ± 0.8 | 35.5 ± 1 | 140.4 ± 0.9 |
| Q4 | 2017 | 505 | 0.95 ± 0.07 | 14.7 ± 2.9 | 39.2 ± 1.8 | 35.6 ± 1 | 140.5 ± 0.8 |
| Q4 | 2018 | 323 | 0.94 ± 0.07 | 13.6 ± 1.5 | 40 ± 0.4 | 35.8 ± 0.5 | 140.9 ± 0.6 |

Table S5. Summary of the top three piecewise structural equation models (SEMs) and full models. Abbreviations are as follows: $K_n$, relative condition factor; TC, habitat temperature; $N_m$, chub mackerel abundance; $N_s$, Japanese sardine abundance; $A$, age; SFOI, the southernmost position of the first Oyashio intrusion; Lat, catch latitude; Lon, catch longitude.

| Quarter | Model | Response variable | Predictor variables | Fisher's $C$ | *p-value* | BIC | ΔBIC |
|---|---|---|---|---|---|---|---|
| Q1 | Full | $K_n$ | $N_m$ + SFOI + $A$ + TC + Lat | 0.00 | 1.00 | 125.00 | 11.41 |
| | | TC | $N_m$ + SFOI + $A$ + Lat | | | | |
| | Best | $K_n$ | $N_m$ + $A$ + TC | 4.21 | 0.38 | 113.58 | 0.00 |
| | | TC | $N_m$ + SFOI + $A$ + Lat | | | | |
| | 2nd | $K_n$ | $N_m$ + $A$ + TC + Lat | 0.88 | 0.64 | 118.07 | 4.49 |
| | | TC | $N_m$ + SFOI + $A$ + Lat | | | | |
| | 3rd | $K_n$ | $N_m$ + SFOI + $A$ + TC | 3.03 | 0.22 | 120.22 | 6.64 |
| | | TC | $N_m$ + SFOI + $A$ + Lat | | | | |
| Q2 | Full | $K_n$ | $N_m$ + $N_s$ + SFOI + $A$ + TC + Lat + Lon | 0.00 | 1.00 | 139.45 | 33.79 |
| | | TC | $N_m$ + $N_s$ + SFOI + $A$ + Lat + Lon | | | | |
| | Best | $K_n$ | $N_m$ + $N_s$ + TC + Lat | 1.07 | 0.98 | 105.66 | 0.00 |
| | | TC | $N_m$ + $A$ + Lat + Lon | | | | |
| | 2nd | $K_n$ | $N_m$ + $N_s$ + Lat | 8.29 | 0.41 | 105.90 | 0.24 |
| | | TC | $N_m$ + $A$ + Lat + Lon | | | | |

| | | | | | | | |
|---|---|---|---|---|---|---|---|
| | 3rd | $K_n$ | $N_m$ + $N_s$ + Lat | 7.98 | 0.24 | 112.57 | 6.91 |
| | | TC | $N_m$ + $N_s$ + $A$ + Lat + Lon | | | | |
| Q3 | Full | $K_n$ | $N_m$ + $N_s$ + SFOI + $A$ + TC + Lat + Lon | 0.00 | 1.00 | 155.40 | 10.15 |
| | | TC | $N_m$ + $N_s$ + SFOI + $A$ + Lat + Lon | | | | |
| | Best | $K_n$ | $N_m$ + SFOI + $A$ + TC + Lat | 5.39 | 0.25 | 145.25 | 0.00 |
| | | TC | $N_m$ + $N_s$ + SFOI + $A$ + Lat + Lon | | | | |
| | 2nd | $K_n$ | $N_m$ + $N_s$ + SFOI + $A$ + TC + Lat | 2.42 | 0.30 | 150.05 | 4.80 |
| | | TC | $N_m$ + $N_s$ + SFOI + $A$ + Lat + Lon | | | | |
| | 3rd | $K_n$ | $N_m$ + SFOI + $A$ + TC + Lat + Lon | 2.97 | 0.23 | 150.61 | 5.36 |
| | | TC | $N_m$ + $N_s$ + SFOI + $A$ + Lat + Lon | | | | |
| Q4 | Full | $K_n$ | $N_m$ + $N_s$ + SFOI + $A$ + TC + Lat | 0.00 | 1.00 | 140.99 | 9.53 |
| | | TC | $N_m$ + $N_s$ + SFOI + $A$ + Lat | | | | |
| | Best | $K_n$ | $N_s$+ SFOI + $A$ + TC | 6.14 | 0.19 | 131.46 | 0.00 |
| | | TC | $N_m$ + $N_s$ + SFOI + $A$ + Lat | | | | |
| | 2nd | $K_n$ | $N_s$ + SFOI + $A$ + TC + Lat | 1.40 | 0.50 | 134.56 | 3.10 |
| | | TC | $N_m$ + $N_s$ + SFOI + $A$ + Lat | | | | |
| | 3rd | $K_n$ | $N_m$ + $N_s$ + SFOT + $A$ + TC | 6.39 | 0.04 | 139.55 | 8.09 |
| | | TC | $N_m$ + $N_s$ + SFOI + $A$ + Lat | | | | |

Table S6. Summary and variance inflation factors (VIFs) of the final model of piecewise structural equation models (SEMs). Abbreviations are as follows: $K_n$, relative condition factor; TC, habitat temperature; $N_m$, chub mackerel abundance; $N_s$, Japanese sardine abundance; $A$, age; SFOI, the southernmost position of the first Oyashio intrusion; Lat, catch latitude; Lon, catch longitude.

| Quarter | Response variable | Predictor | Estimate | SE | *p*-value | Standardized estimate | VIF |
|---|---|---|---|---|---|---|---|
| Q1 | $K_n$ | $N_m$ | −0.0034 | 0.0007 | <0.001 | −0.255 | 1.26 |
| | | $A$ | −0.0137 | 0.0012 | <0.001 | −0.265 | 1.04 |
| | | TC | 0.0012 | 0.0014 | 0.382 | 0.046 | 1.23 |
| | TC | $N_m$ | −0.2034 | 0.0100 | <0.001 | −0.414 | 1.49 |
| | | SFOI | 0.2821 | 0.0738 | <0.001 | 0.065 | 1.07 |
| | | $A$ | 0.3438 | 0.0392 | <0.001 | 0.179 | 1.48 |
| | | Lat | −0.9121 | 0.0600 | <0.001 | −0.290 | 1.31 |
| Q2 | $K_n$ | $N_m$ | −0.0036 | 0.0017 | 0.042 | −0.237 | 2.49 |
| | | $N_s$ | −0.0019 | 0.0008 | 0.029 | −0.254 | 2.40 |
| | | TC | −0.0018 | 0.0029 | 0.544 | −0.053 | 1.39 |
| | | Lat | 0.0200 | 0.0040 | <0.001 | 0.335 | 1.50 |
| | TC | $N_m$ | 0.0679 | 0.0115 | <0.001 | 0.151 | 1.10 |
| | | $A$ | −0.2526 | 0.0733 | <0.001 | −0.092 | 1.18 |
| | | Lat | −0.4710 | 0.0514 | <0.001 | −0.266 | 1.35 |

| | | | | | | | |
|---|---|---|---|---|---|---|---|
| | | Lon | −1.2294 | 0.0877 | <0.001 | −0.409 | 1.45 |
| Q3 | $K_n$ | $N_m$ | −0.0048 | 0.0008 | <0.001 | −0.281 | 1.43 |
| | | SFOI | −0.0310 | 0.0067 | <0.001 | −0.255 | 1.52 |
| | | $A$ | 0.0187 | 0.0019 | <0.001 | 0.236 | 1.05 |
| | | TC | −0.0054 | 0.0023 | 0.021 | −0.121 | 1.63 |
| | | Lat | 0.0070 | 0.0029 | 0.018 | 0.129 | 1.51 |
| | TC | $N_m$ | −0.1418 | 0.0093 | <0.001 | −0.366 | 2.37 |
| | | $N_s$ | 0.0178 | 0.0042 | <0.001 | 0.109 | 2.68 |
| | | SFOI | −1.0508 | 0.0495 | <0.001 | −0.385 | 1.35 |
| | | $A$ | −0.1901 | 0.0346 | <0.001 | −0.107 | 1.50 |
| | | Lat | −0.1118 | 0.0385 | 0.004 | −0.092 | 3.97 |
| | | Lon | −1.1237 | 0.0948 | <0.001 | −0.353 | 3.48 |
| Q4 | $K_n$ | $N_s$ | −0.0032 | 0.0003 | <0.001 | −0.485 | 1.24 |
| | | SFOI | 0.0093 | 0.0029 | 0.002 | 0.140 | 1.09 |
| | | $A$ | 0.0162 | 0.0015 | <0.001 | 0.212 | 1.09 |
| | | TC | 0.0072 | 0.0014 | <0.001 | 0.213 | 1.13 |
| | TC | $N_m$ | 0.0669 | 0.0115 | <0.001 | 0.141 | 3.27 |
| | | $N_s$ | −0.0739 | 0.0049 | <0.001 | −0.380 | 3.58 |
| | | SFOI | 0.5678 | 0.0283 | <0.001 | 0.290 | 1.16 |
| | | $A$ | 0.3408 | 0.0356 | <0.001 | 0.151 | 1.38 |
| | | Lat | −1.7931 | 0.0399 | <0.001 | −0.669 | 1.24 |

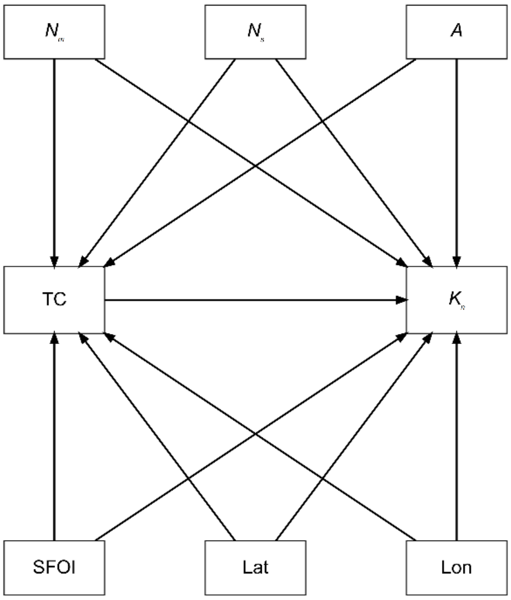


Figure S1. *A priori* structural equation model of relationships among relative condition factor ($K_n$), chub mackerel abundance ($N_m$), Japanese sardine abundance ($N_s$), age ($A$), habitat temperature (TC), the southernmost position of the first Oyashio intrusion (SFOI), catch latitude (Lat), and catch longitude (Lon).

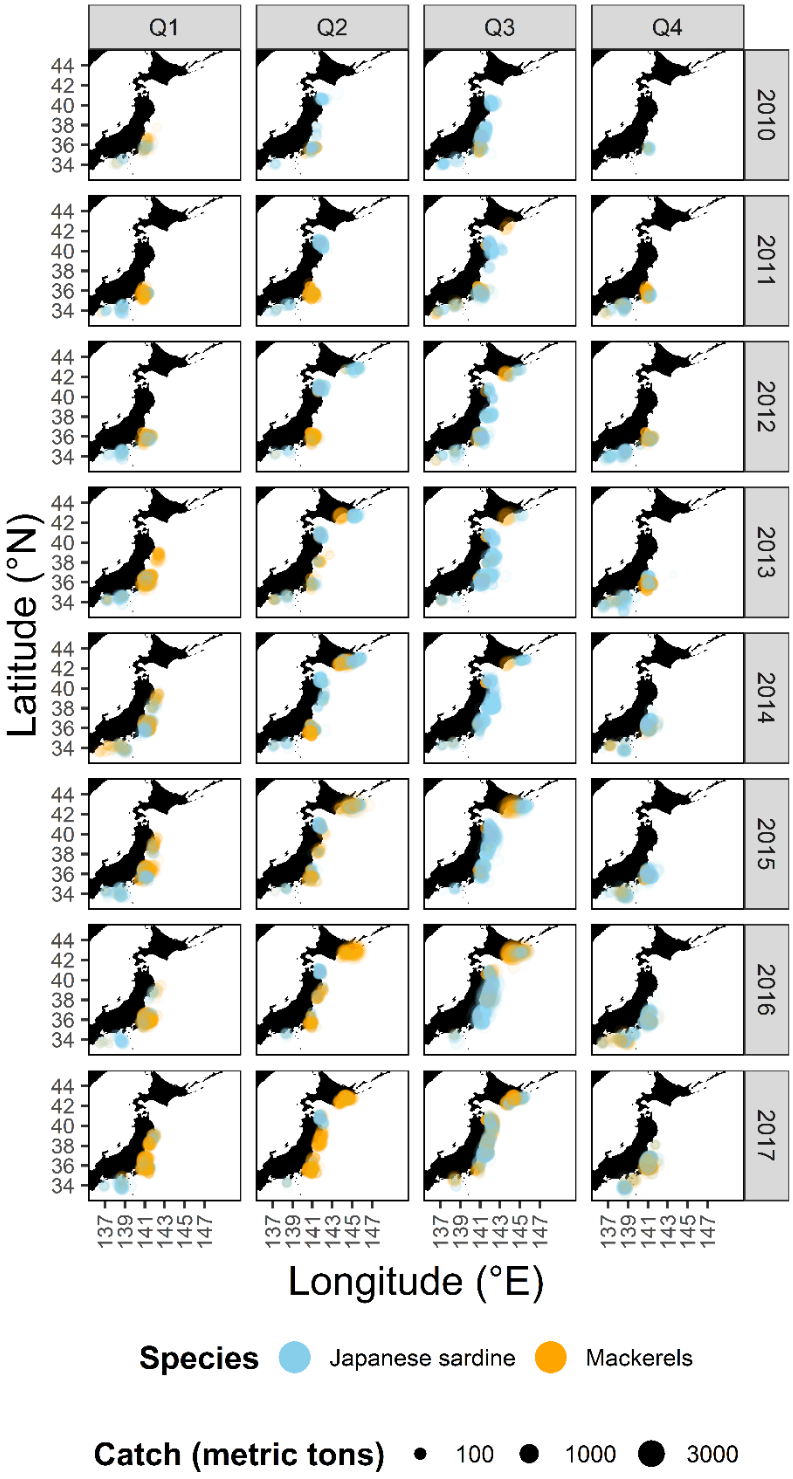


Figure S2. Seasonal changes in the catch locations of mackerels (figure shows combined catches of chub and spotted mackerel) and Japanese sardine by purse seine fisheries from 2010 to 2017.

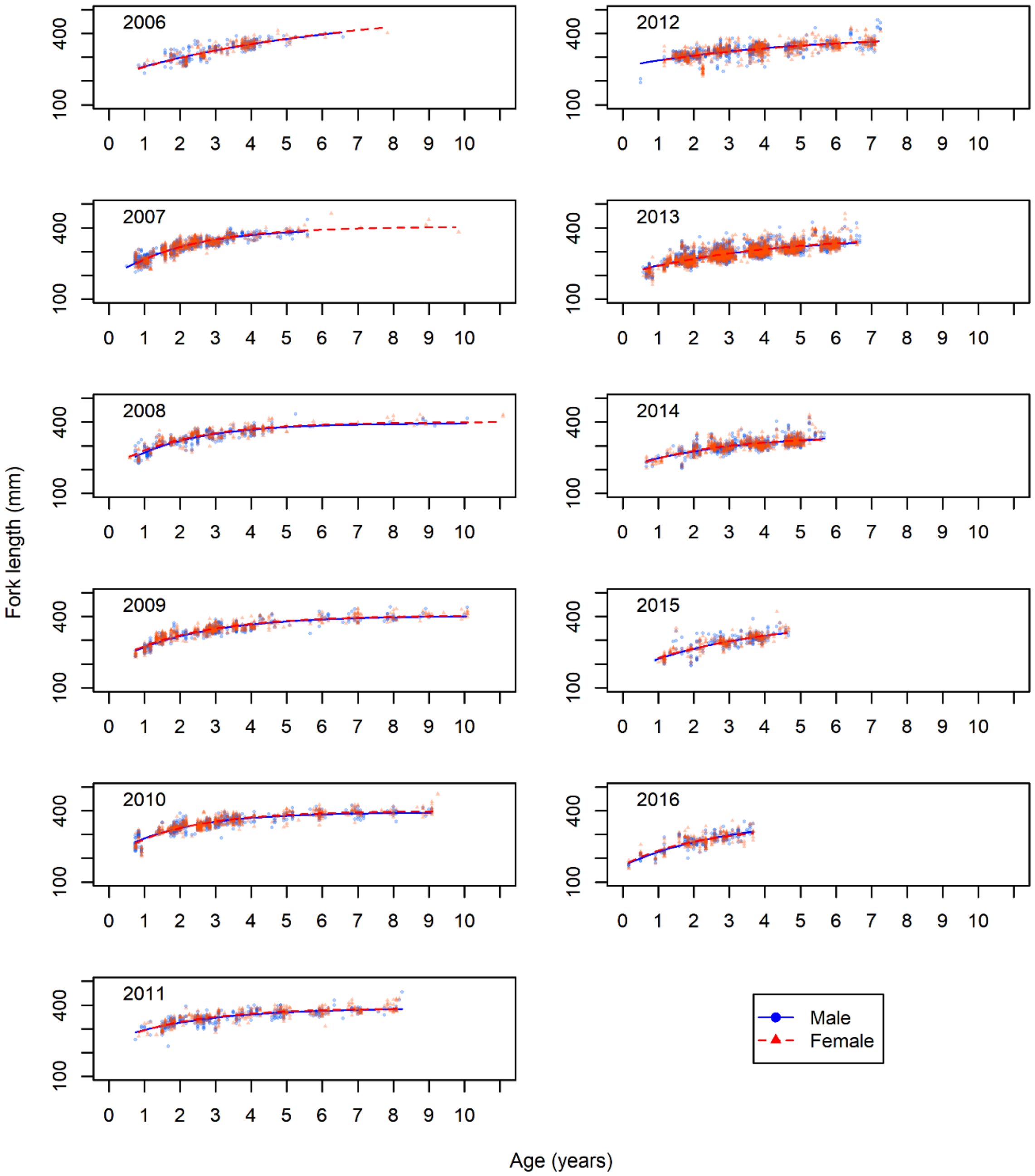


Figure S3. Comparison of lengths-at-age estimated by using von Bertalanffy growth formulae among sexes and year classes. Red dashed lines and triangles show data for females, and blue solid lines and circles show data for males.

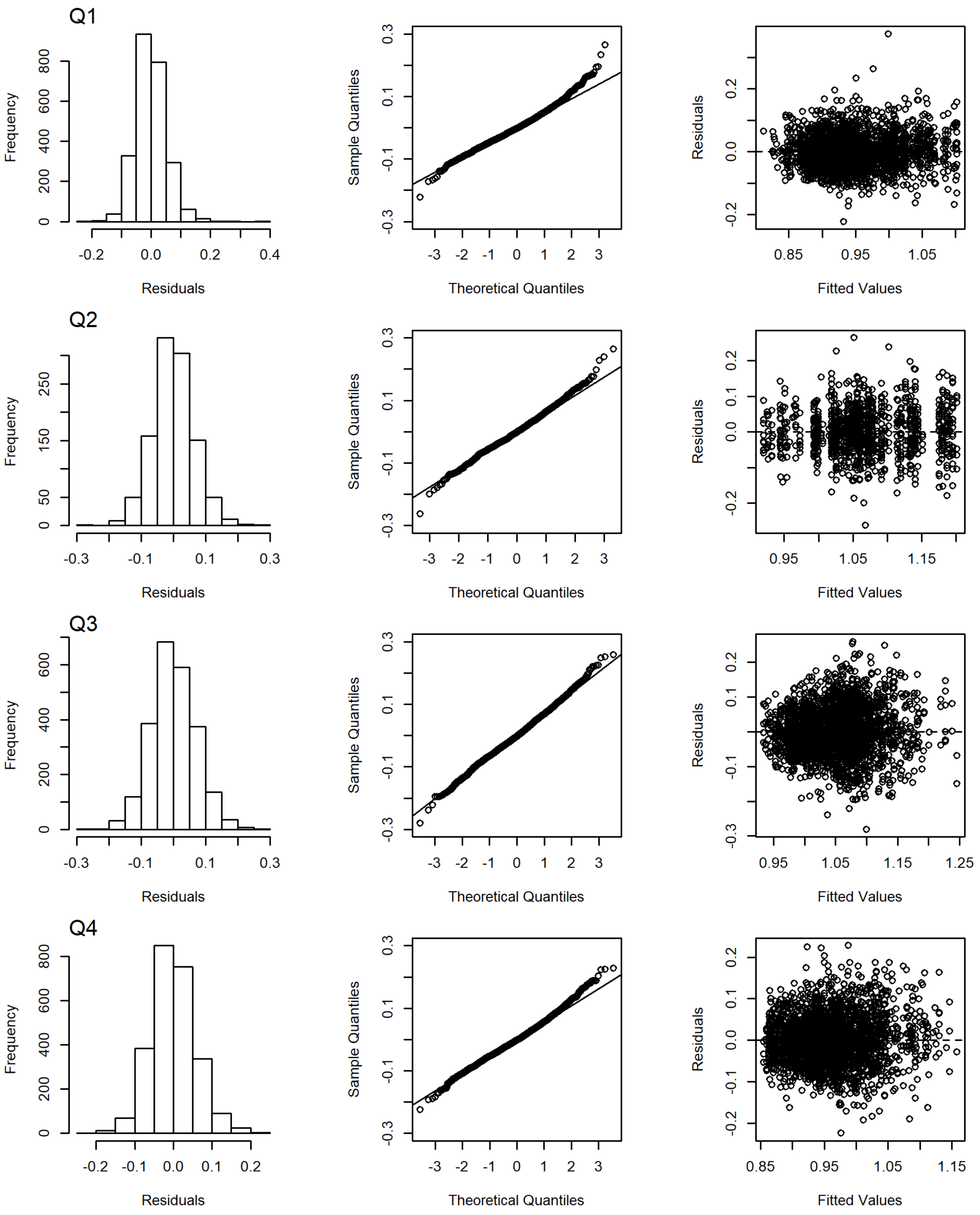


Figure S4. Histograms of the distribution of residuals, Q–Q plots, and residuals vs. fitted values for the $K_n$ model by quarter.

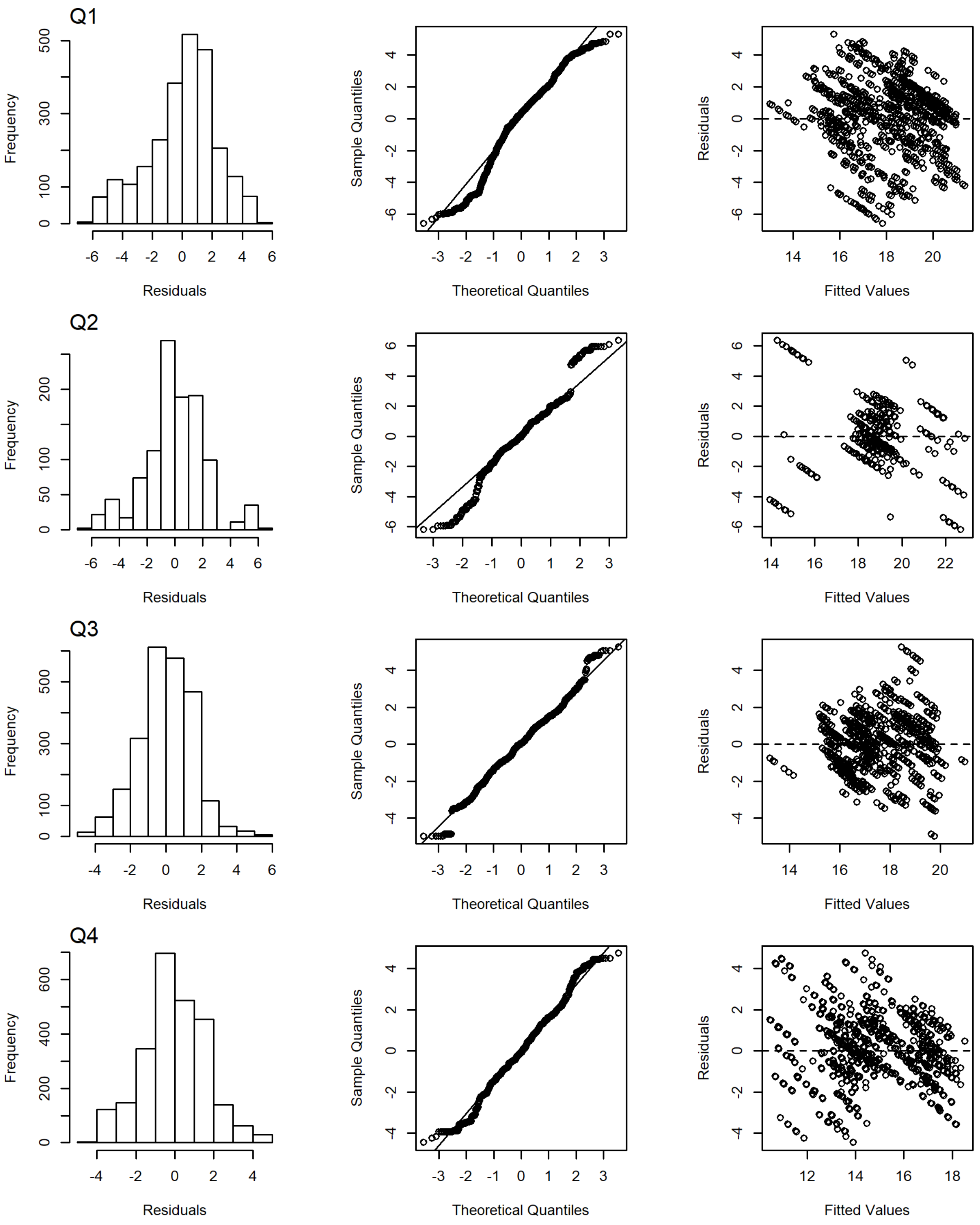


Figure S5. Histograms of the distribution of residuals, Q–Q plots, and residuals vs. fitted values for the TC model by quarter.
